# Cortical contraction drives the 3D patterning of epithelial cell surfaces

**DOI:** 10.1101/819987

**Authors:** Aaron P. van Loon, Ivan S. Erofeev, Ivan V. Maryshev, Andrew B. Goryachev, Alvaro Sagasti

**Affiliations:** Department of Molecular, Cell and Developmental Biology and Molecular Biology Interdepartmental Program, University of California, Los Angeles; Centre for Synthetic and Systems Biology, School of Biological Sciences, University of Edinburgh

**Author notes:** Authors contributed equally (Aaron P. van Loon, Ivan S. Erofeev).

## Abstract

Cellular protrusions create complex cell surface topographies, but biomechanical mechanisms regulating their formation and arrangement are largely unknown. To study how protrusions form, we focused on the morphogenesis of microridges, elongated actin-based structures projecting from the apical surfaces of zebrafish skin cells that are arranged in labyrinthine patterns. Microridges form by accreting simple finger-like precursors. Live imaging demonstrated that microridge morphogenesis is linked to apical constriction. A non-muscle myosin II (NMII) reporter revealed pulsatile contractions of the actomyosin cortex; inhibiting NMII demonstrated that contractions are required for apical constriction and microridge formation. A biomechanical model suggested that contraction reduces surface tension to permit the fusion of precursors into microridges. Indeed, reducing surface tension with hyperosmolar media promoted microridge formation. In anisotropically stretched cells, microridges formed by precursor fusion along the stretch axis, which computational modeling explained as a consequence of stretch-induced cortical flow. Collectively, our results demonstrate how contraction within the 2D plane of the cortex patterns 3D cell surfaces.

**SUMMARY:** Microridges, elongated 3D protrusions arranged in maze-like patterns on zebrafish skin cells, form by the accretion of simple precursor projections. Modeling and in vivo experiments showed that cortical contractions promote the coalescence of precursors into microridges by reducing membrane tension.

## INTRODUCTION

Animal cells generate a broad repertoire of dynamic structures based on the highly versatile and plastic actin cytoskeleton (Pollard and Cooper, 2009; Blanchoin et al., 2014). Actin generates both the protrusive forces that shape the membrane and, in conjunction with myosin, contractile forces that can alter cell geometry. Rapid restructuring of the actin cytoskeleton is controlled by a core of conserved actin regulatory proteins, including nucleators, elongators, bundlers, depolymerizers, and myosin motors (Pollard, 2016). Despite their universality, the divergent patterns of self-organization between these regulators generate a remarkable diversity of actin-based structures, including filopodia, lamellipodia, microvilli, dorsal ruffles, and podosomes (Blanchoin et al., 2014; Buccione et al., 2004). While actin regulatory proteins have been extensively studied, neither molecular mechanisms, nor biophysical principles that generate and switch between specific actin structures are well understood. The coexistence and competition of distinct actin-based structures within the same cell makes these problems even more complex (Rotty and Bear, 2014; Lomakin et al., 2015).

Microridges are membrane protrusions extended in one spatial dimension and arranged in remarkable, fingerprint-like patterns on the apical surface of mucosal epithelial cells (Fig. 1A) (Straus, 1963; Olson and Fromm, 1973). Microridges are found in a wide array of species on a variety of tissues, including the cornea, oral mucosa, and esophagus (Depasquale, 2018), and are thought to aid in mucus retention (Sperry and Wassersug, 1976; Pinto et al., 2019). Microridges are filled with actin filaments and associate with several actin-binding proteins (Depasquale, 2018; Pinto et al., 2019). Interestingly, microridges do not emerge as fully spatially-extended structures like dorsal ruffles. Instead, they assemble from short vertically-projecting precursors (Raman et al., 2016; Lam et al., 2015; Uehara et al., 1988; Gorelik et al., 2003). Ultrastructural analyses have demonstrated that actin filaments in microridges have mostly branched actin networks (Bereiter-Hahn et al., 1979; Pinto et al., 2019) and, therefore, it is unclear if microridge precursors are more similar in their actin organization to podosomes or microvilli, to which they had been frequently compared. To emphasize this distinction, we have dubbed these precursors actin “pegs”. Inhibiting Arp2/3 prevents aggregation of actin pegs into microridges, suggesting that branched actin networks are also required for microridge assembly (Lam et al., 2015; Pinto et al., 2019). Factors regulating non-muscle myosin II (NMII) activity have been found to promote microridge elongation (Raman et al., 2016), but reports differ about whether NMII plays a direct role in microridge morphogenesis (Lam et al., 2015).

**Figure 1.**
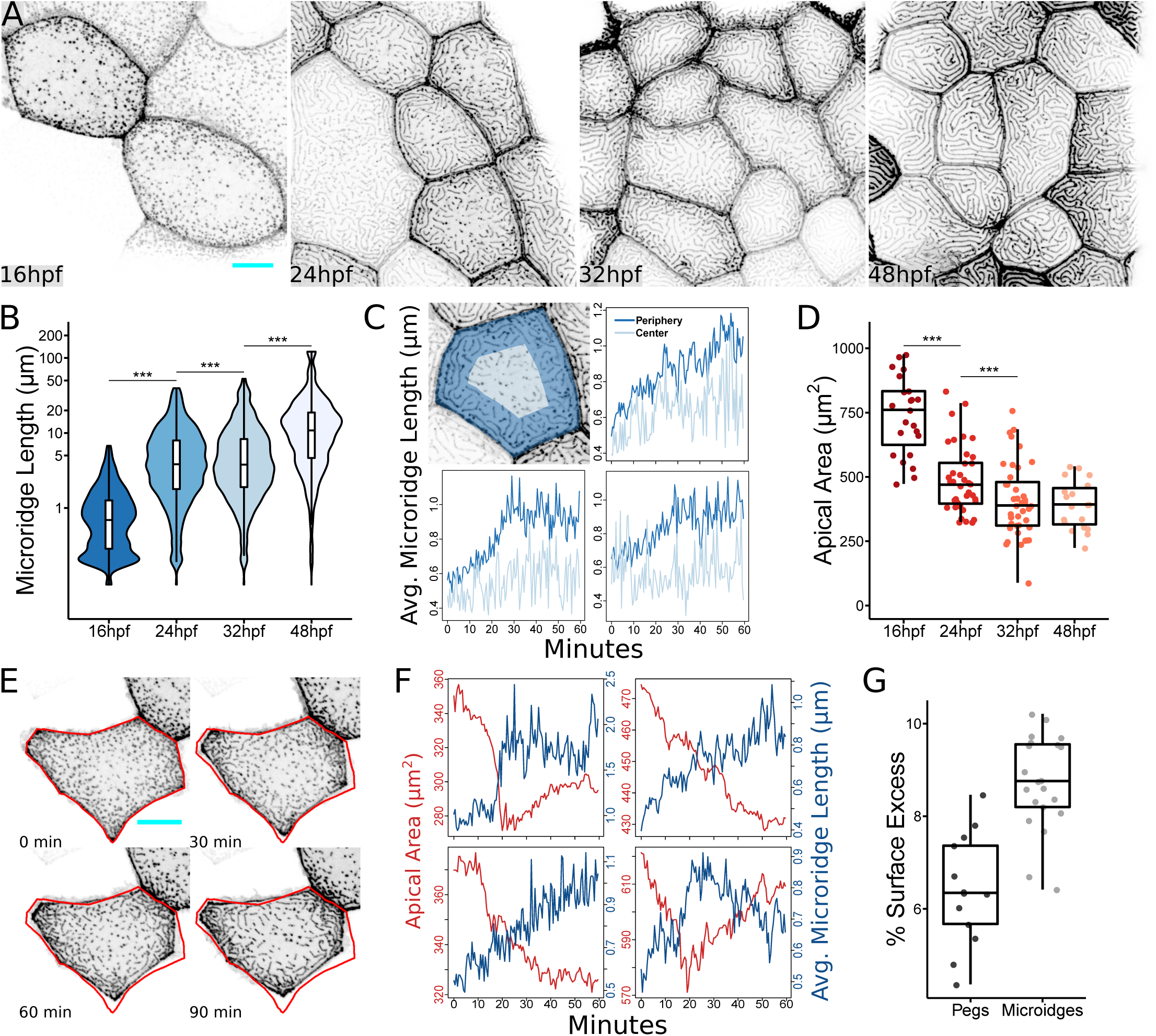
Microridge length changes in tandem with apical cell area (A) Representative projections of Lifeact-GFP in periderm cells on zebrafish larvae at the indicated stages of zebrafish development. (B) Box and violin plot of microridge length at the indicated stages of zebrafish development. Data displayed is a weighted distribution of microridge length, in which frequency is proportional to microridge length, approximating occupied area. For a non-weighted presentation of the same data see Fig. S1K. ***p < 0.0001; ANOVA followed by Tukey’s HSD test (n=15582 structures in 23 cells from 10 fish at 16 hpf; n=5096 structures in 40 cells from 9 fish at 24 hpf; n=4572 structures in 40 cells from 9 fish at 32 hpf; n=1309 structures in 19 cells from 6 fish at 48 hpf). (C) Upper left panel: Diagram of cell “periphery” (dark blue) and “center” (light blue) zones, representing 75% and 25% of apical cell area, respectively. Other panels: Line graphs comparing the average microridge length in the cell “periphery” versus the cell “center” over time. (D) Dot and box plot of periderm cell apical area at the indicated stages of zebrafish development. *p < 0.01; ***p < 0.0001; ANOVA followed by Tukey’s HSD test (n=23 cells from 10 fish at 16 hpf; n=40 cells from 9 fish at 24 hpf; n=40 cells from 9 fish at 32 hpf; n=19 cells from 6 fish at 48 hpf). (E) Sequential projections from a time-lapse movie of Lifeact-GFP in a single periderm cell during apical constriction. Red outline: position of cell border at 0 min. (F) Line plots of apical area and average microridge length in single periderm cells over time. Upper right panel corresponds to cell shown in (E). (G) Dot and box plot of surface excess (relative difference between total surface area and projected surface area as seen in microscope) in regions of the apical cell membrane composed of only microridges or only pegs. (n=13 regions with pegs, n=21 regions with microridges). Further details of this analysis are provided in Materials and Methods. Scale bars: 10μm (A and E)

Although microridges are less studied than other actin-based structures, they offer an excellent opportunity to probe systemic properties of cytoskeletal regulation. Microridge patterns possess several characteristic parameters, including their spatial orientation, length distribution, and periodicity, which can be readily quantified from live images. These parameters reflect biochemical and biomechanical processes that regulate the morphogenesis of actin structures and are sensitive to experimental intervention into these processes. Multiple genetic and pharmacological perturbations can thus be applied to dissect principles of patterning and test theoretical hypotheses.

By imaging microridge development on the skin of larval zebrafish, we have found that cortical contraction couples apical constriction to microridge morphogenesis. In vivo experiments and modeling suggest that contraction of the apical actomyosin cortex relieves surface tension to facilitate the coalescence of pegs to form, elongate, and orient microridges. Thus, cortical contraction not only determines the size and shape of the apical surface, but also concomitantly sculpts its 3D surface.

## RESULTS

### The apical surfaces of periderm cells shrink as microridges form

We first asked if we could identify overarching organizational principles in the emergence of microridges from pegs. To characterize the process of microridge development in live animals, we imaged transgenic zebrafish expressing the F-actin reporter Lifeact-GFP specifically in periderm cells during development (Rasmussen et al., 2015; Helker et al., 2013). We developed an automated image analysis protocol to segment microridges from these images and quantify microridge length in an unbiased manner (Fig. S1). As previously reported (Lam et al., 2015; Raman et al., 2016; Pinto et al., 2019), early in development (16 hours post-fertilization, hpf), periderm cells projected actin pegs that superficially resemble short microvilli (Fig. 1A-B). By 24 hpf, elongated microridges appeared near cell borders, whereas pegs still populated the center of the apical cell surface (Fig. 1A-B). By 32 hpf, microridges filled the apical surface, and continued to elongate through at least 48 hpf (Fig. 1A-B). This temporal progression of microridge growth was apparent from plotting the distribution of the pooled population of protrusions (Fig. 1B, S2A), or from measuring the average protrusion length per cell (Fig. S2B). To determine how cells transitioned from pegs to microridges, we imaged microridge growth in live animals at 15-30-second intervals. These videos revealed that pegs were dynamic, and coalesced to both form and elongate microridges (Fig. S3, Video 1). Time-lapse imaging also demonstrated that microridges form in a centripetal manner: assembly of microridges from pegs started in the cell periphery and progressed towards the cell center (Fig. 1C, Video 1). These observations confirmed previous studies suggesting that actin pegs are precursors to microridges that coalesce to form and elongate microridges (Lam et al., 2015; Pinto et al., 2019).

We next considered if microridge morphogenesis is associated with other changes in the morphology or biomechanical properties of the developing epithelium. Indeed, we noticed that during the period of transition from pegs to microridges (∼16-32 hpf) the apical area of periderm cells decreased (44.7% on average), but stabilized between 32 hpf and 48 hpf (Fig. 1D). Moreover, average microridge length in individual cells inversely correlated with apical cell area: smaller cells had, on average, longer microridges (Fig. S2C), suggesting that apical area may influence microridge length. To determine whether cell areas shrunk predominantly by apical constriction or cell division, we imaged actin dynamics at 30-second intervals during an early stage of microridge elongation (18-19 hpf). These videos demonstrated that cells underwent intermittent bouts of apical constriction and relaxation, but predominantly constricted, similar to the ratchet-like process that has been described in other instances of apical constriction (Martin et al., 2009; Solon et al., 2009; Blanchard et al., 2010). Microridge length closely tracked changes in apical cell area: microridges elongated, likely by peg accretion, as apical areas shrunk, and microridges shortened as apical areas increased (Fig. 1E-F). We conclude that pegs and microridges are in a dynamic equilibrium and that apical constriction promotes microridge formation.

### A model for microridge formation reproduces experimental observations

Apical constriction significantly reduces the 2D-projected apical area of epithelial cells, as illustrated by our live-cell imaging (Fig. 1E-F). However, it was not clear how actin pegs and microridges, which determine the 3D topography of the membrane, affect the total surface area of the apical membrane. We therefore asked whether cells with pegs or cells with microridges have a larger 3D apical surface when their projected apical areas are identical. To answer this question, we assumed that actin pegs and microridges are of equal and uniform height and computationally evaluated the total 3D surface of apical regions with only microridges or only pegs. Surprisingly, after normalization by the 2D-projected area of the regions, we found that microridges induce larger membrane surfaces (Fig. 1G). Therefore, while apical constriction reduces the apical membrane surface, the associated transition from pegs to microridges increases it.

To gain quantitative insight into these observations, we developed a simple biophysical model of the cellular apical domain. We hypothesized that apical morphogenesis is generated by the dynamics of three closely interacting subsystems with distinct biomechanical properties: the membrane itself, the immediately underlying branched actin structure that fills pegs and microridges, and the deeper actomyosin cortex (Fig. 2A). Actin filaments within the branched structure are largely disordered but project into the neighboring membrane (Uehara et al., 1991; Pinto et al., 2019), similar to the actin structures that power lamellipodia protrusion. We thus assumed that their polymerization stretches the membrane and expands the membrane surface. Conversely, filaments in the deeper actomyosin cortex are largely aligned parallel to the basal surface of cells (Pinto et al., 2019). Contraction of the cortex drives apical constriction (Martin and Goldstein, 2014) and reduces membrane tension (Fig. 2B). Excess membrane is presumably removed by endocytosis (Sonal et al., 2014). We thus propose that the two actin subsystems have opposing effects on membrane area and tension; branched actin and the contractile actomyosin cortex increase and decrease membrane tension, respectively. Pattern formation in our model is driven by the autocatalytic branched polymerization of actin at the membrane-cytoskeleton interface. To ensure formation of both pegs and microridges, we resorted to a prototypical activator-inhibitor model of Gierer and Meinhardt (Gierer and Meinhardt, 1972), in which the role of the inhibitor is played by the height *h* of the actin structure. This heuristic assumption mimics the opposition that membrane tension produces to actin polymerization (Gov, 2006; Gov and Gopinathan, 2006; Atilgan et al., 2006; Mogilner and Rubinstein, 2005). Conversely, membrane tension is relaxed by myosin motor-driven contraction of the actomyosin cortex, whose dynamics are described by the well-established active gel model (Prost et al., 2015). A detailed description of the model equations and parameters are provided in the Methods.

**Figure 2.**
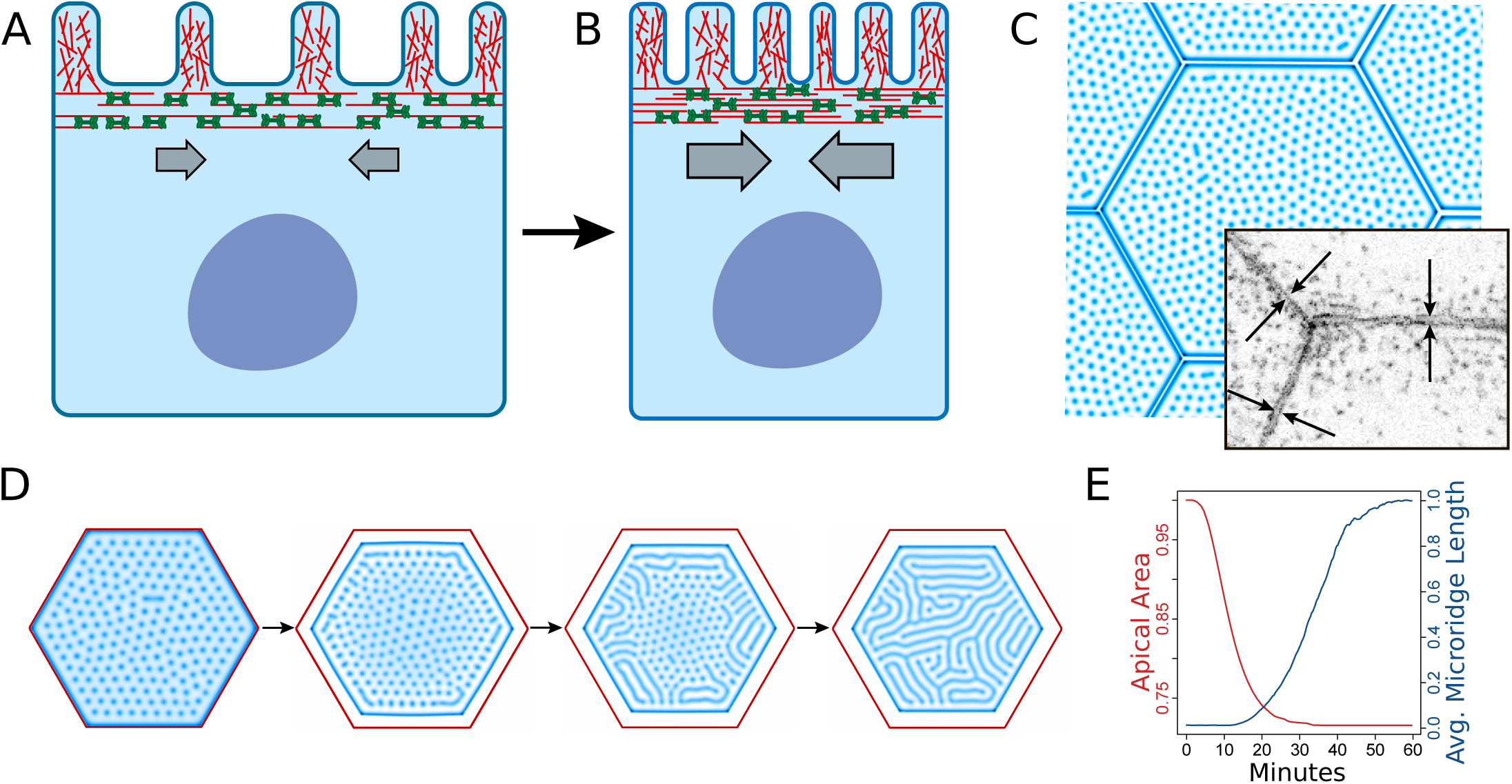
*In silico* simulation of apical constriction mimics microridge development *in vivo* (A) Diagram of a periderm cell in homeostatic conditions with actin-filled microridges projecting from the apical surface. The underlying apical cortex is rich in actin (red filament) and NMII (green mini-filament) and attached to the cell membrane. (B) Diagram of a periderm cell undergoing apical constriction. NMII contraction in the apical cortex relieves tension in the attached cell membrane, allowing actin to protrude. (C) Cells from *in silico* simulations developed a long microridge at the cell border prior to microridge formation elsewhere on the apical membrane. Arrows in the inset micrograph point to a similar structure in periderm cells expressing Lifeact-GFP prior to microridge development. (D) *In silico* simulation of apical constriction in our biomechanical model recapitulates the centripetal progression of microridge development observed *in vivo*.

In our model, spatially uniform isotropic contraction of the apical actomyosin cortex produced a transition from pegs to microridges, which occurred uniformly on the apical surface (Video 2). We noticed that even with parameters corresponding to the initially relaxed cortex, when the interior of the apical domain is populated only by pegs, a single closed microridge had formed immediately proximal to the cell boundary (Fig. 2C). Interestingly, such microridges positioned next to the tight junctions between cells have been routinely observed in experiments by us and others (Depasquale, 2018). Remarkably, as in the model, they typically form prior to the formation of microridges in the interior of the apical domain. In live-cell images, they emerged first as discontinuous, paired structures positioned on each side of, and strictly parallel to, the tight junctions (Fig. 2C, inset). As microridges developed within the apical interior, these junction-associated structures matured into proper microridges and continuously surrounded the entire apical domains of cells (Fig. 1A). In the model, formation of this outer microridge is determined by the boundary condition that fixes vertical membrane deflection on the boundary to *h = 0*. Thus, our model predicts that formation of these circumferential microridges is determined by the singularity in the membrane tension imposed by unyielding tight junctions.

Model simulations showed that uniform contraction of the apical actomyosin cortex cannot explain centripetal emergence of ridges from the cell boundaries. We thus surmised that the apical actomyosin cortex could also undergo a process of maturation. Possibly, its contractility increases first at the tight junctions, where Rho GTPase activity that drives actin polymerization and myosin contraction are typically enriched (Zihni and Terry, 2015; Ratheesh et al., 2012), and then progresses inwards. Simulations of the model augmented with this additional hypothesis reproduced the experimental observations. Starting at the cell boundary, pegs coalesced into microridges, which eventually filled the entire apical domain (Fig. 2D-E, Video 2).

### NMII is required for apical constriction and microridge formation

A previous study suggested that NMII is involved in lengthening microridges (Raman et al., 2016), but did not determine how it contributes to microridge formation, nor whether it is linked to apical constriction. We therefore sought direct evidence that NMII produces apical constriction and induces microridge morphogenesis from actin pegs. We first inhibited NMII contractility by treating zebrafish larvae with the specific small molecule inhibitor blebbistatin (Straight et al., 2003) for 24 hr spanning the period of microridge development (16-40 hpf). Blebbistatin reduced apical constriction in a concentration-dependent manner and inhibited the coalescence of pegs into microridges (Fig. 3A-C). Since extended exposures to blebbistatin could affect microridges in a variety of direct or indirect ways, we examined the effects of shorter treatments; periderm cells expressing the actin reporter were imaged before and after 2 hr of blebbistatin exposure. During this short period of exposure, blebbistatin significantly inhibited microridge elongation and modestly reduced apical constriction, compared to controls (Fig. 3E-F). In control cells, microridge length and apical cell area were inversely correlated (R^2^ = −.65), but this relationship was diminished by treatment with blebbistatin (R^2^ = −.31) (Fig. 3G).

**Figure 3.**
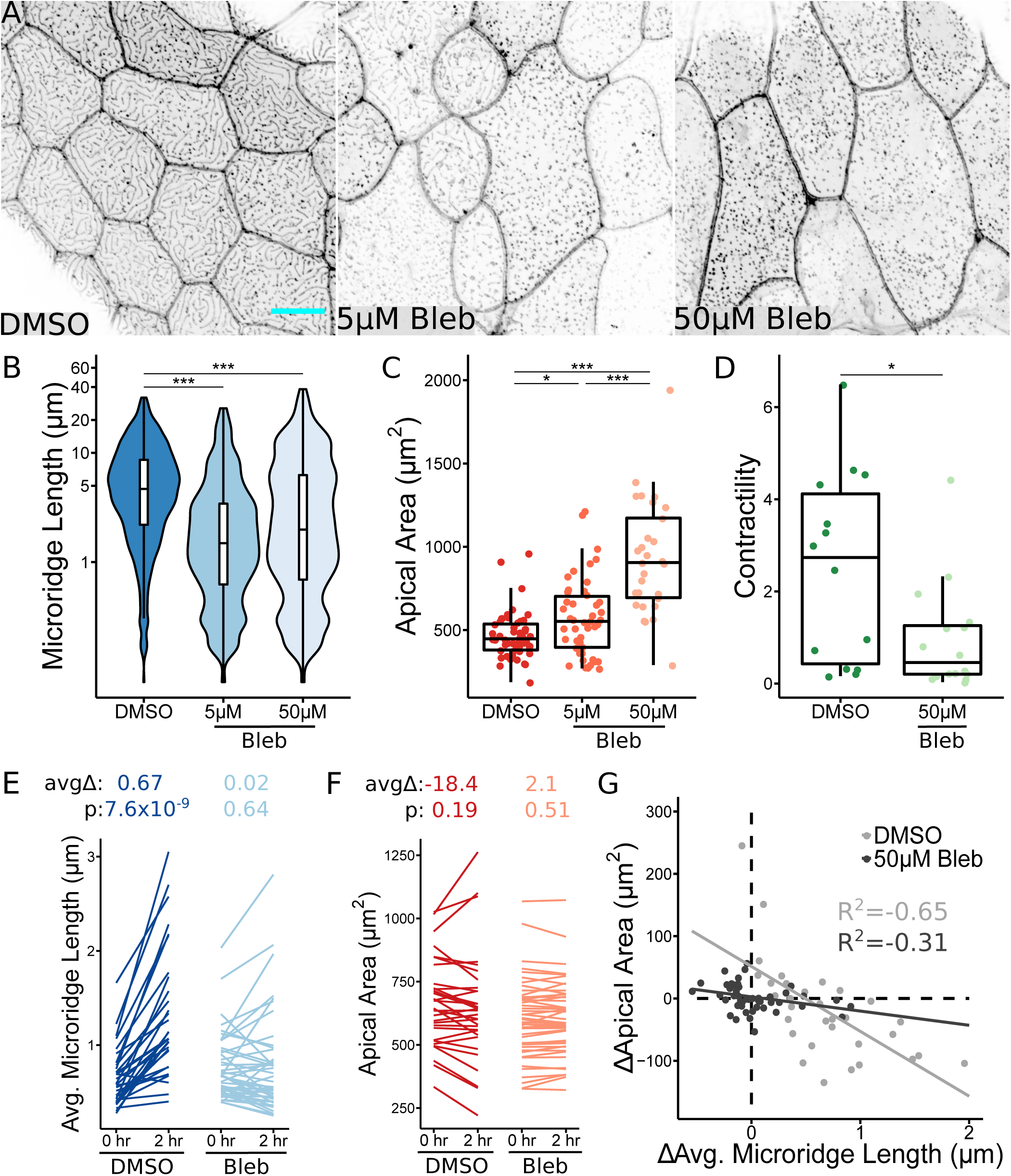
Non-muscle Myosin II contraction is required for apical constriction and microridge development (A) Representative projections of Lifeact-GFP in periderm cells on 40 hpf zebrafish larvae after 24 hr treatment with either 1% DMSO or indicated concentration of blebbistatin. (B) Box and violin plot of microridge length in 40 hpf zebrafish embryos after 24 hr treatment with either 1% DMSO or indicated concentration of blebbistatin. Data is presented as a weighted distribution of microridge length, in which frequency is proportional to length, approximating occupied area. ***p < 0.0001; ANOVA followed by Tukey’s HSD test (n=6772 structures in 53 cells from 10 fish for 1% DMSO, n=9587 structures in 46 cells from 11 fish for 5μM Blebbistatin; n=8623 structures in 29 cells from 13 fish for 50μM Blebbistatin). (C) Dot and box plot of periderm cell apical area in 40 hpf zebrafish embryos after 24 hr treatment with either 1% DMSO or indicated concentration of blebbistatin. *p < 0.01; ***p < 0.0001; ANOVA followed by Tukey’s HSD test (n=53 cells from 10 fish for 1% DMSO, n=46 cells from 11 fish for 5μM Blebbistatin, n=29 cells from 13 fish for 50μM Blebbistatin). (D) Dot and box plot of total NMII reporter contraction area summed over a 10-minute period after 1 hr treatment with either 1% DMSO or 50μM blebbistatin. *p < 0.01; student’s t-test (n=10416 contractions in 29 cells from 18 fish for 1% DMSO, n=2259 contractions in 16 cells from 9 fish for 50μM Blebbistatin). (E) Line plots of average microridge length in individual periderm cells before (18 hpf) and after (20 hpf) 2 hr treatment with either 1% DMSO or 50μM Blebbistatin. Above, average change in average microridge length and p values (paired t-test). (n=17039 structures in 64 cells from 10 fish for 1% DMSO, n=20873 structures in 92 cells from 11 fish for 50μM Blebbistatin). (F) Line plots of periderm cell apical area in individual cells before (18 hpf) and after (20 hpf) 2 hr treatment with either 1% DMSO or 50μM Blebbistatin. Above, average change in periderm cell apical area and p values (paired t-test). (n=64 cells from 10 fish for 1% DMSO, n=92 cells from 11 fish for 50μM Blebbistatin). (G) Scatter plot of change in average microridge length versus change in periderm cell apical area after 2 hr treatment with 1% DMSO or 50μM Blebbistatin. R^2^ determined using Pearson’s correlation coefficient. (n=17039 structures in 64 cells from 10 fish for 1% DMSO, n=20873 structures in 92 cells from 11 fish for 50μM Blebbistatin). Scale bars: 10μm (A)

The branched actin nucleator Arp2/3 is required for microridge formation and maintenance (Lam et al., 2015). As expected, an inhibitor of Arp2/3, CK666 (Nolen et al., 2009), prevented coalescence of actin pegs into microridges, but did not prevent pegs from forming or reduce their dynamics (Fig. S4, and not shown). Interestingly, however, CK666 failed to reduce apical constriction and, in fact, significantly promoted it (Fig. S4C). This observation is consistent with our hypothesis that polymerization of the branched actin subsystem, for which Arp2/3 is required, induces membrane surface expansion and, thus, opposes apical constriction, which is driven by the underlying actomyosin layer.

To directly visualize the localization and activity of NMII in periderm cells during apical constriction and microridge development, we created a transgenic zebrafish line that expresses an NMII reporter (Maître et al., 2012) specifically in periderm cells. As expected, this reporter localized to cell-cell junctions and appeared to be distributed across the apical cortex. Time-lapse imaging revealed transient, local flashes of reporter fluorescence at the apical surface, which we interpreted as contractile pulses that concentrated NMII at their foci (Fig. 4A, Video 3). Indeed, one hour of exposure to blebbistatin was sufficient to significantly decrease these pulses (Fig. 3D), confirming that they reflect contractile activity of NMII. These apical NMII pulses temporally and spatially resembled pulsatile contractions that drive apical constriction in other systems (Fernández et al., 2007; Solon et al., 2009; Blanchard et al., 2010; David et al., 2010). Contractile events concentrated towards the periphery of zebrafish periderm cells early in microridge development (16 hpf), progressed towards the center as development proceeded (24 hpf), and dampened after microridge formation (48 hpf) (Fig. 4B), supporting our hypothesis that apical constriction initiates at, and centripetally propagates from, junctions. This outside-in progression of cortical activity mirrors the spatial progression of microridge formation (Fig. 1C). Time-lapse imaging of periderm cells expressing both actin and NMII reporters demonstrated that contraction events pulled nearby actin pegs towards myosin foci (Fig. 4C-D, Video 3).

**Figure 4.**
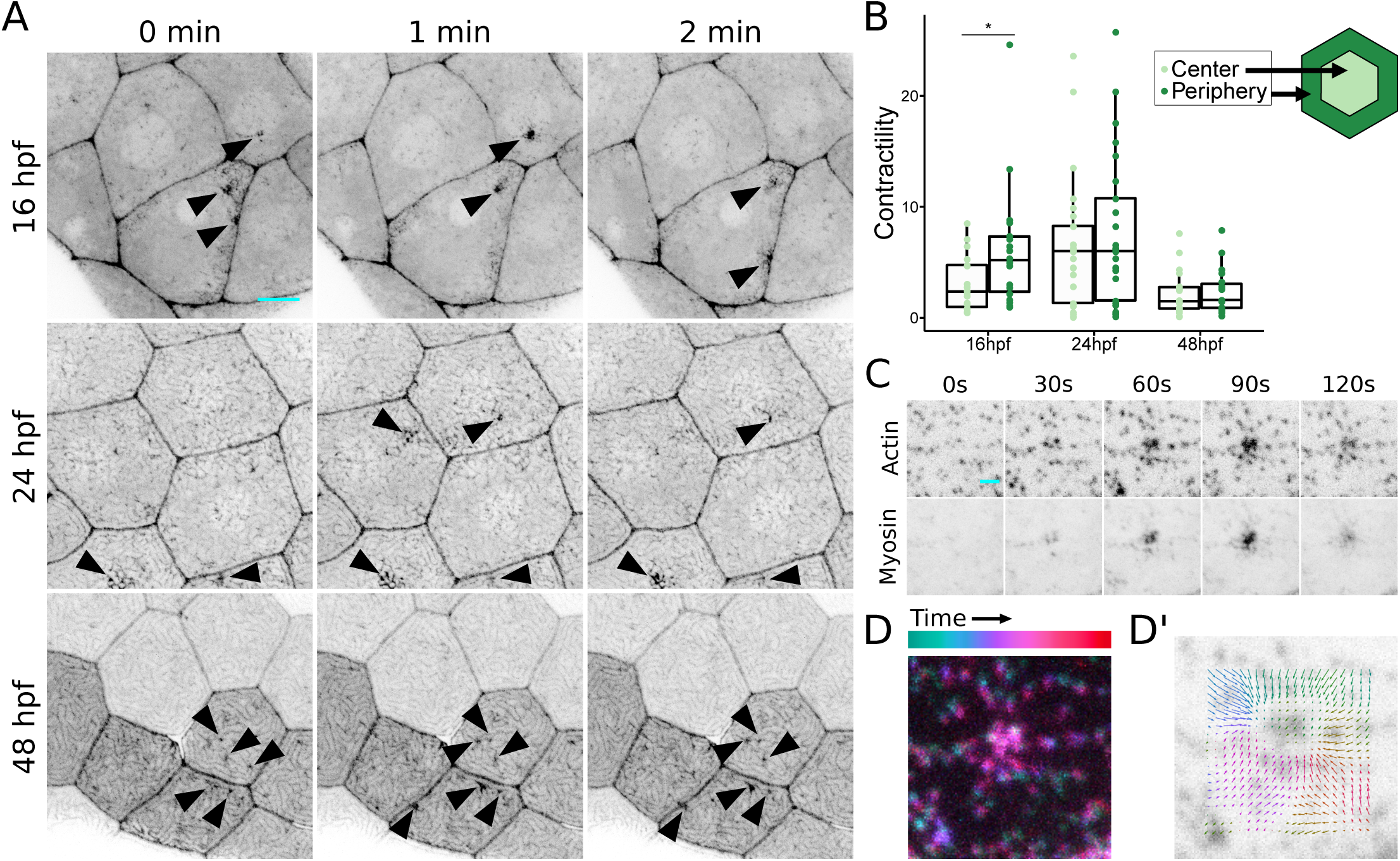
Apical Non-muscle Myosin II contractions pinch the cell membrane (A) Sequential projections from time-lapse movies of Myl12.1-EGFP in periderm cells at indicated stages of zebrafish development. Arrowheads: dynamic concentrations of NMII reporter fluorescence at the apical membrane. (B) Dot and box plot of total NMII reporter contraction area summed over a 10-minute period at specified time points during zebrafish development. Contractions were categorized based on whether the majority of contraction area was inside the inner 25% of the cell surface (Center) or in the remaining outer 75% (Periphery). *p < 0.01; student’s t-test (n=11794 contractions in 19 cells from 13 fish at 16hpf; n=18776 contractions in 25 cells from 13 fish at 24 hpf; n=6303 contractions in 19 cells from 7 fish at 48 hpf). (C) Sequential projections from a time-lapse movie of Lifeact-Ruby and Myl12.1-EGFP during an apical NMII pulse in a 16 hpf periderm cell. (D-D’) Superimposition of sequential frames from a time-lapse movie, and particle image velocimetry (PIV) (D’) show the centripetal trajectory of actin structures towards the focus of contraction. Scale bars: 10μm (A) and 2μm (C)

Contractile activity of NMII is activated via phosphorylation of the regulatory myosin light chain (MLC) by multiple kinases, such as Rho GTPase effector kinase (ROCK) and myosin light chain kinase (MLCK) (Matsumura, 2005). To determine if these kinases regulate microridge morphogenesis, we inhibited ROCK or MLCK with the small molecule inhibitors Rockout or ML-7, respectively, between 16-24 hpf. While ML-7 had no effect on microridge formation (data not shown), Rockout significantly decreased microridge length and increased apical cell area in a concentration-dependent manner (Fig. 5A-C). Additionally, one hour of Rockout treatment reduced NMII pulses (Fig. 5D). Rockout did not dramatically affect peg dynamics, indicating that contraction specifically regulates peg coalescence, not peg formation (data not shown). This result indirectly supports the hypothesis that contraction of the actomyosin cortex is driven by activity of RhoA via its effector ROCK. We conclude that, regardless of its upstream regulation, NMII-driven contraction of the apical actomyosin cortex is required for both apical constriction and formation of microridges from actin peg precursors.

**Figure 5.**
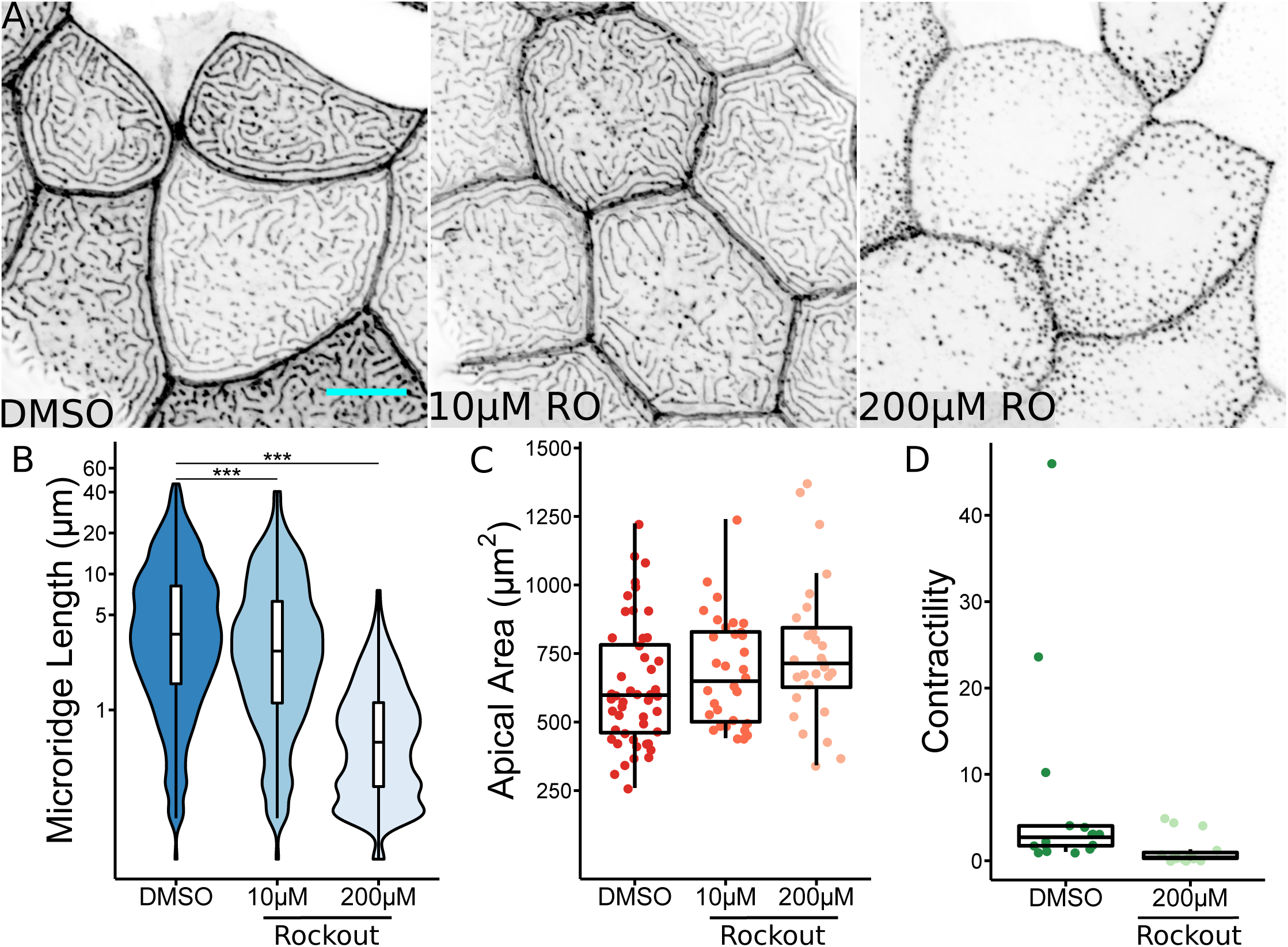
ROCK activity is required for microridge development (A) Representative projections of Lifeact-GFP in periderm cells on 24 hpf zebrafish larvae after 8 hr treatment with either 0.2% DMSO or indicated concentration of Rockout. (B) Box and violin plot of microridge length in 24 hpf zebrafish embryos after 8 hr treatment with either 0.2% DMSO or indicated concentration of Rockout. Data displayed is a weighted distribution of microridge length where frequency is proportional to microridge length, approximating occupied area. ***p < 0.0001; ANOVA followed by Tukey’s HSD test (n=10385 structures in 49 cells from 15 fish for 0.2% DMSO, n=8353 structures in 34 cells from 11 fish for 10μM Rockout, n=7501 structures in 28 cells from 10 fish for 200μM Rockout). (C) Dot and box plot of periderm cell apical area in 24 hpf zebrafish embryos after 8 hr treatment with either 0.2% DMSO or indicated concentration of Rockout. p = 0.074; ANOVA (n=49 cells from 15 fish for 0.2% DMSO, n=34 cells from 11 fish for 10μM Rockout, n=28 cells from 10 fish for 200μM Rockout). (D) Dot and box plot of total NMII reporter contraction area summed over a 10-minute period after 1 hr treatment with either 0.2% DMSO or 200μM Rockout. p = 0.075; student’s t-test (n=9048 contractions in 16 cells from 10 fish for 0.2% DMSO, n=3495 contractions in 16 cells from 10 fish for 200μM Rockout). Scale bars: 10μm (A)

### Membrane tension directly controls microridge formation

Although the activity of NMII and its activation by ROCK are required for apical constriction and microridge formation, it is possible that myosin affects microridge formation by a means not related to its biomechanical function; for example, by serving as a scaffold for signaling complexes. We therefore sought to directly test whether membrane tension or an unrelated function of NMII controls microridge formation. To alter membrane tension, we exposed zebrafish embryos to hyperosmolar media during early stages of microridge development. Placing animals in a hyperosmolar environment should draw water from periderm cells, causing them to “deflate”, thus reducing membrane tension independent of myosin contraction. Indeed, exposing zebrafish embryos to either high salt media or glycerol-supplemented media at a stage when cells are dominated by actin pegs, but before significant microridge formation typically occurs (16 hpf), caused cells to shrink rapidly. Time-lapse imaging demonstrated that as cells shrank, actin pegs rapidly coalesced into microridges (Fig. 6, Video 4). Thus, reducing membrane tension is sufficient to promote microridge formation, in agreement with our *in silico* model.

**Figure 6.**
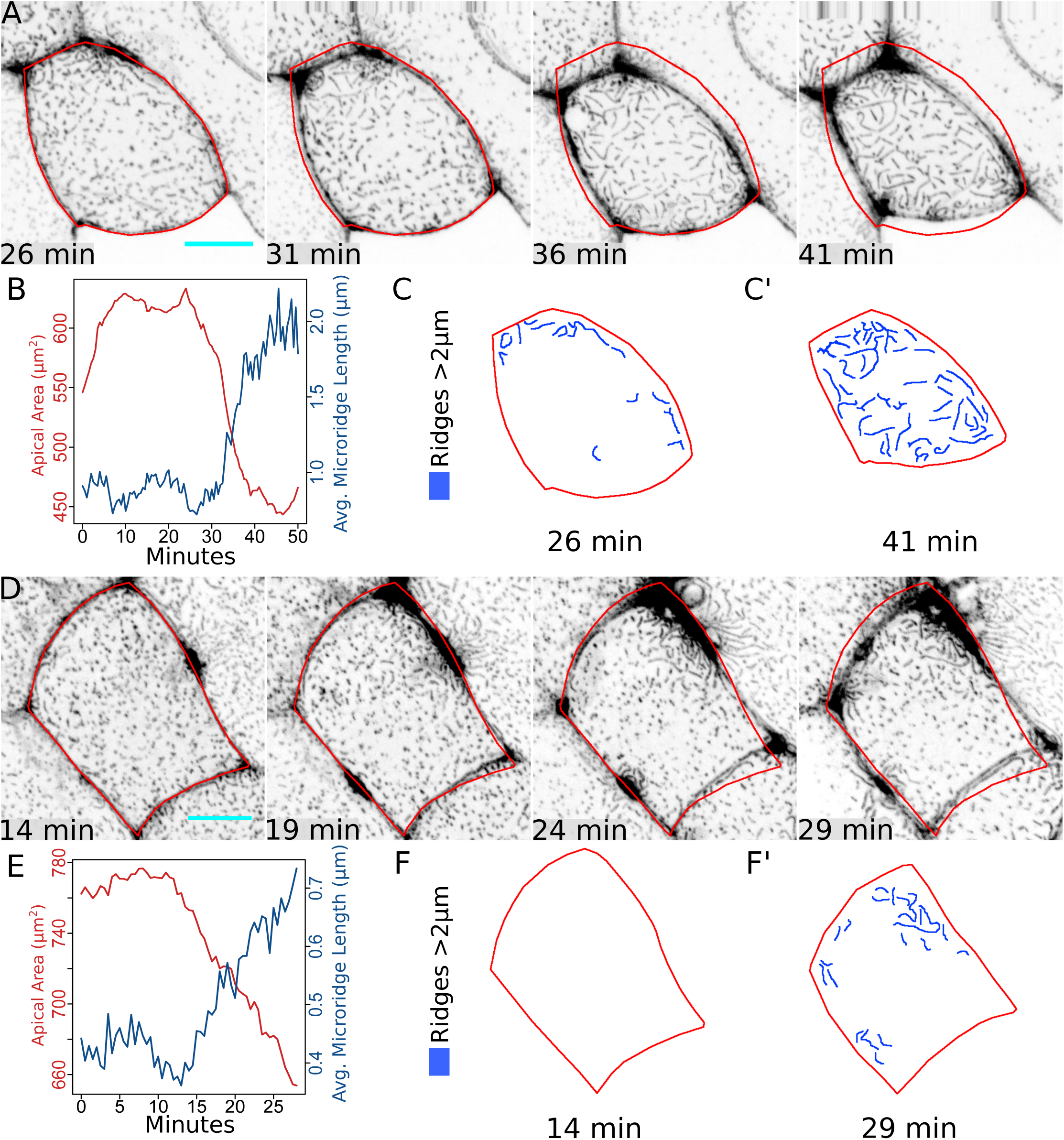
Membrane surface energy regulates microridge formation (A) Sequential projections from a time-lapse movie of Lifeact-GFP in periderm cells after exposure to water with 500x Instant Ocean salt. Red outline: position of cell border at 26 min. (B) Line plot of apical area and average microridge length in the periderm cell shown in A after exposure to water with 500x Instant Ocean salt. (C-C’) Diagram of cell in A at the indicated time points with microridges longer than 2μm traced in blue. (D) Sequential projections from a time-lapse movie of Lifeact-GFP in periderm cells after exposure to 12.5% glycerol. Red outline: position of cell border at 14 min. (E) Line plot of apical area and average microridge length in the cell shown in D after exposure to 12.5% glycerol. (F-F’) Diagram of cell in E at the indicated time points with microridges longer than 2μm traced in blue. Scale bars: 10μm (A and D)

### Anisotropy of microridge formation indicates that peg coalescence is an active process

To further our understanding of microridge formation, we sought to direct this process in a well-controlled experimental set-up. To achieve this, we leveraged the natural wound-healing behavior of epithelial sheets. In response to ablation of individual cells, neighboring cells generate a powerful biomechanical response to rapidly constrict the wound (Lam et al., 2015; Rosenblatt et al., 2001). If two cells are ablated simultaneously, intervening cells will sometimes undergo near perfect uniaxial stretching along the axis connecting the two wounds (Fig. 7A-C). For these experiments, we chose to ablate periderm cells at an early developmental stage with few microridges (16 hpf), using a laser on a 2-photon microscope (Video 5). Cells were selected for analysis if they exhibited a robust uniaxial stretch. These analyses showed that stretch was accompanied by a small apical surface reduction, on average 10% of the projected area (Fig. 7E). Remarkably, as they stretched, all cells formed new microridges, which were predominantly aligned along the stretch axis (Fig. 7B-D, F, Video 5). The highest anisotropy of microridge orientation was coincident with the maximal distortion of stretched cells (∼10 min after ablation). This alignment slightly decreased as the epithelium relaxed into a new steady-state configuration.

**Figure 7.**
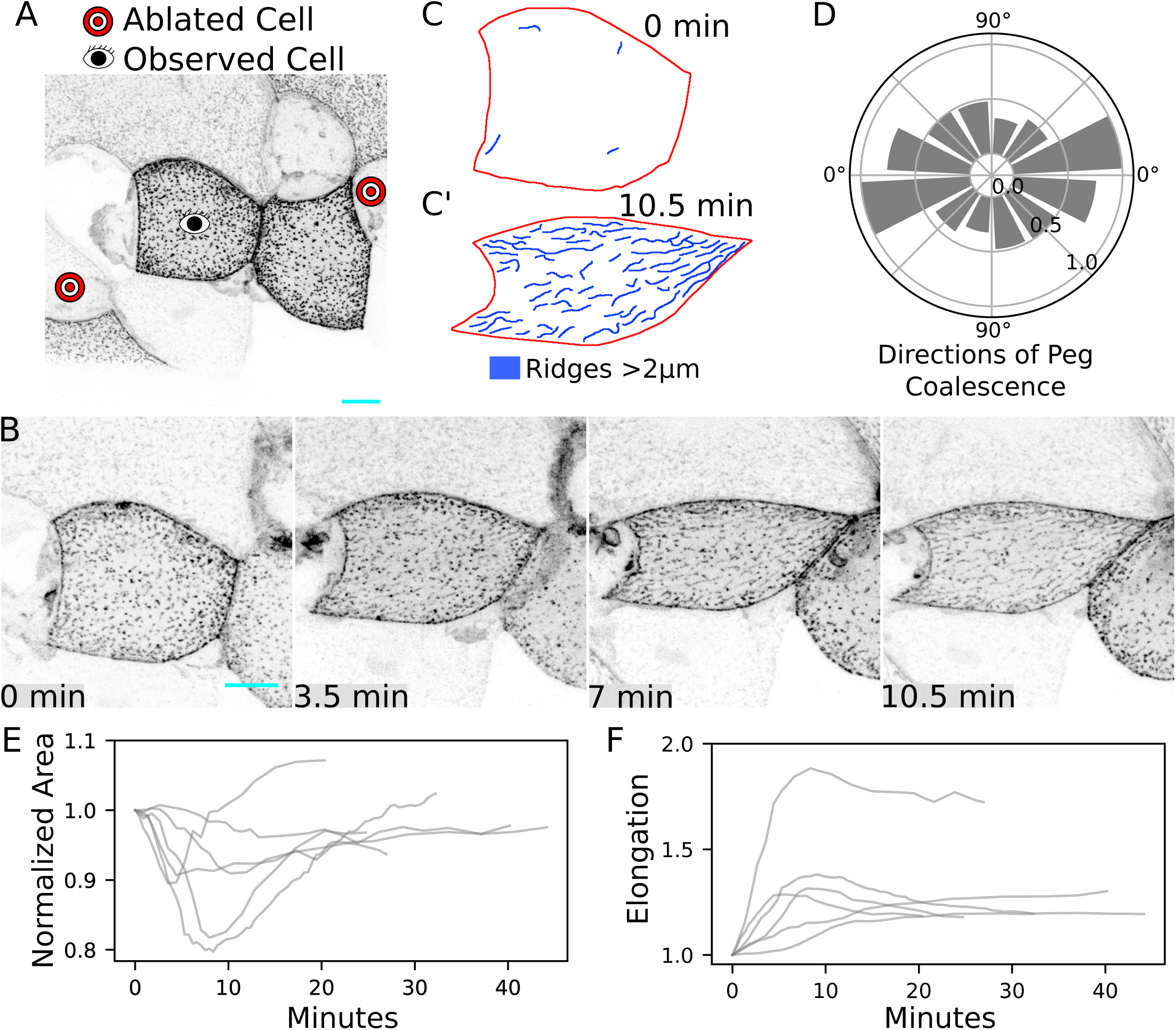
Cortical organization and diffusion direct microridge formation and orientation (A) Projection of Lifeact-GFP in periderm cells on a 16 hpf zebrafish embryo before laser cell ablation. Eye: observed cell. Target: cell to be ablated. (B) Still images from a time-lapse sequence of a cell elongating over 10 minutes. (C) Outline of the elongating cell in B showing the initial time-point (C), and 10 minutes after ablation (C’). Microridges longer than 2 µm are highlighted blue. (D) Averaged histogram of directions of peg coalescence events (normalized to maximal bin, direction of elongation at 0°). (E) Projected area for 5 cells used in this analysis. Areas were normalized to the initial value. (F) Elongation factor (ratio of longest axis to shortest axis for the transformation of the cell) for 5 analyzed cells. Scale bars: 10μm (A and B)

Cell stretch produced by neighbor ablation temporarily induces a flow of the viscoelastic actomyosin cortex, which is transmitted to the plasma membrane and the underlying branched F-actin cortex via multiple protein-protein links. The observed orientation of microridges along the stretch axis could be potentially explained by two distinct sources, both induced by flow. First, the torque generated by the actomyosin flow could re-orient preexisting microridges along the direction of stretch. However, quantification (Fig. 7C,C’) showed that, prior to cell stretch, microridges were essentially nonexistent and largely formed during the stretch itself. Thus, reorientation of preexisting microridges contributes little, if at all, to the aligned microridges seen in the experiment. Alternatively, microridges could form in an oriented manner if the fusion of pegs occurred preferentially along the direction of stretch. To test this second hypothesis, we quantified the angle at which actin peg fusion occurred after laser ablation. This analysis demonstrated that, in all analyzed cells, peg fusion was strongly anisotropic, on average three times more frequent along the stretch axis than perpendicular to it (Fig. 7D), confirming that microridges indeed form in an oriented manner.

The observation that pegs fuse along the stretch axis is surprising, as actin pegs sandwiched between the membrane and the actomyosin cortex are transported by the cortical flow and, thus, would be expected to collide preferentially along the direction orthogonal to the stretch axis (Fig. 8A). Theory predicts that peg fusion is energetically preferable (Gov, 2006; Derényi et al., 2002), as it reduces membrane bending energy. Hydrodynamic flow-induced collision of pegs should reduce the potential barrier to fusion and, therefore, promote peg fusion perpendicular to the direction of stretch. Indeed, in agreement with this theoretical argument, and contrary to experimental results, simulations of our model that emulated cell stretch produced preferential fusion of pegs perpendicular, rather than parallel, to the stretch axis (Fig. 8C-D). This discrepancy suggested that our model failed to capture the full complexity of cortical biomechanics. Hydrodynamic flow could potentially order initially isotropic actin filaments along the stretch axis and, thus, induce orientation of force-generating NMII filaments (Fig. 8B). Counterintuitively, this passive re-orientation would increase the active stress generated by the actomyosin gel in the direction of stretch and reduce it in the opposite direction. Introduction of this hypothesis into our model produced simulation results in full agreement with the experiment (Fig. 8C, E-F, Video 6). Furthermore, the model revealed the existence of a minimal value of the flow-induced actomyosin anisotropy, below which peg fusion still occurs predominantly perpendicular to the stretch axis (Fig. 8F). Since our analysis of experimental data identified a significant (3:1) preference for pegs to fuse along the stretch axis, we conclude that stretch-induced cortical flow must induce substantial orientation of F-actin fibers and NMII motors.

**Figure 8.**
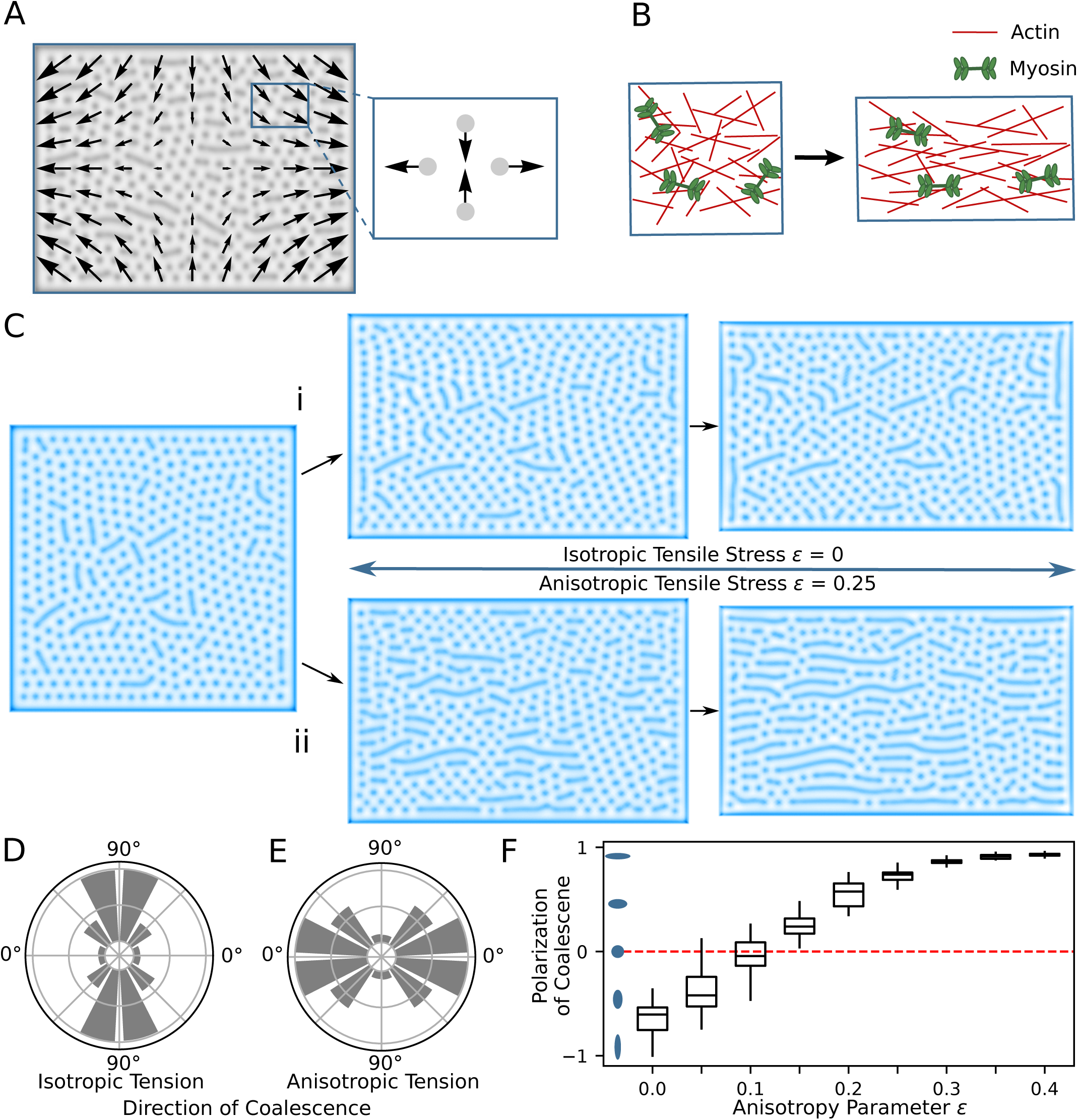
Flow-induced actomyosin anisotropy directs peg fusion along the cell stretch axis (A) Approximated cortex flow during uniaxial stretch. Flow can be locally represented as elongation along one axis and compression along an orthogonal axis. (B) Scheme of rapid stretch that leads to partial alignment of actin and myosin bundles. (C) Modelling of cell stretching with (i) isotropic and (ii) anisotropic tensile stress. (D, E) Corresponding histograms of distributions of structure fusions (averaged over 100 simulations with different initial conditions). If tension does not depend on the anisotropic flow, the fusions happen mostly in the direction perpendicular to the elongation (D). In the case of anisotropic tensile stress, the fusions happen in the direction of elongation (E). (F) Polarization of coalescence histogram for different values of the anisotropy parameter. Polarization value −1 corresponds to coalescence in the perpendicular direction, value 1 corresponds to coalescence in the direction parallel to elongation.

## DISCUSSION

NMII-based contraction is well known to alter cell surfaces in two dimensions: for example, polarized contractions at junctions regulate polarized cell rearrangements (Bertet et al., 2004; Blankenship et al., 2006), and cortical contraction shrinks surfaces during apical constriction (Martin and Goldstein, 2014). Here we demonstrate that contraction in zebrafish periderm cells not only changes 2D cell surface geometry, but also simultaneously sculpts the 3D topography of cells: NMII-based cortical contractions couple apical constriction to the patterning of microridges, which protrude from the apical surface of zebrafish periderm cells, orthogonal to the cortex. Computational modeling and in vivo experiments together support a model in which cortical contractions lower membrane surface tension to permit the coalescence of actin pegs into microridges, and cortical flow influences the organization of contractile machinery, which determines microridge orientation.

Microridges form by the accretion of precursor structures (pegs), a feature that distinguishes them from better-studied protrusions, such as lamellipodia and dorsal ruffles, which emerge and expand as a single unit. Inhibiting Arp2/3, NMII, or ROCK prevented peg coalescence into microridges, but did not appear to affect pegs themselves. Thus, peg formation and peg coalescence are two separate morphogenetic steps under distinct biomechanical and biochemical regulation. Understanding peg formation, actin dynamics in pegs, and the regulation of peg density will be critical to fully understanding microridge morphogenesis. For example, aggregation of similar peg-like precursors to form ridge-like structures in cultured kidney cells is influenced by actin dynamics (Gorelik et al., 2003). A critical local peg density may be required for microridge formation, but our observations suggest that it is not sufficient, since peg density remains relatively constant in the peripheral domain of cells for several hours before microridge formation begins (not shown). Upon integrating into microridges, pegs may, at least in part, retain their integrity as substructures since we, and others (Depasquale, 2018), have noted intensely labeled F-actin puncta within microridges. In fact, one ultrastructural study reported that periodic actin bundles could be discerned within microridges by electron microscopy (Bereiter-Hahn et al., 1979), though a more recent study using high resolution techniques did not identify these substructures (Pinto et al., 2019). Whether pegs contain bundled parallel actin filaments, like microvilli, or branched filaments, like podosomes, is thus unclear. Identifying the bundling proteins, nucleators, and motors that localize to pegs may resolve whether they resemble microvilli or podosomes, or are unique structures with a distinct actin organization and protein composition.

The cortical contractions we observed in periderm cells resemble contractions driving well-characterized behaviors in other, better studied systems, like *C. elegans* and *Drosophila* gastrulation (Roh-Johnson et al., 2012; Martin et al., 2009). In those other systems, contraction is driven by Rho family GTPase signaling networks (Mason et al., 2013; Munjal et al., 2015; Marston et al., 2016). It is thus likely that contraction of zebrafish periderm cells during microridge formation also depends on Rho family GTPases. Indeed, we found that the RhoA effector ROCK is required for apical contraction and microridge development in periderm cells, and previous work showed that RhoA inhibition can alter microridge patterning (Lam et al., 2015). Our observation that contractions initially predominate near cell borders may result from the association of RhoA regulators with cell-cell junctions (Ratheesh et al., 2012; Zihni and Terry, 2015), and could contribute to the centripetal progression of micoridge formation. Defining the contribution of cell-cell junctions and Rho signaling networks could help explain how cortical contractions are tuned to create biomechanical conditions conducive to apical constriction and microridge morphogenesis.

Our theoretical model predicted that reducing surface tension is sufficient to promote peg coalescence into microridges. This prediction was supported by experiments showing that microridges rapidly formed as cells shrank in hyperosmolar media. Conversely, cell swelling would be predicted to prevent micoridge formation or cause microridge disassembly. Unfortunately, fish larvae and periderm cells appeared to be unaffected by treatment with hypo-osmolar media (not shown), potentially due to homeostatic regulatory mechanisms. Nonetheless, a previous study showed that in zebrafish with Myosin Vb mutations, cells swell due to defects in vesicular trafficking and lose microridges (Sonal et al., 2014), consistent with the idea that increasing membrane surface tension opposes microridge formation.

Contractile patterns are shaped by the flow of the contractile machinery and NMII regulators within the plane of the cortex (Munjal et al., 2015; Rauzi et al., 2010; Bray and White, 1988). Our experiments and biomechanical modeling demonstrated that when cortical flow is anisotropically oriented by cell stretching, pegs coalesce into microridges in an oriented manner, aligning nascent microridges along the stretch axis. This phenomenon could explain microridge orientation during naturally anisotropic cell behaviors, like cytokinesis. Just before cytokinesis, cells expand and microridges dissolve back into pegs; during cytokinesis, cells contract dramatically and microridges rapidly re-form (Lam et al., 2015). These new microridges are initially oriented predominantly perpendicular to the cytokinetic furrow, consistent with the observation that, during cytokinesis, furrow ingression drives polarized cortical flow towards the furrow (Khaliullin et al., 2018; Cao and Wang, 1990; DeBiasio et al., 1996).

Micoridges (and closely related structures called microplicae) are found on a variety of mucosal tissues in many vertebrate animals, suggesting that they play a common role in mucus retention (Depasquale, 2018). Nonetheless, their morphologies vary significantly in length, spacing, and branching, perhaps reflecting optimized morphologies for their function in different tissue contexts. Intriguingly, microridge morphology even varies in different parts of the zebrafish skin that are likely under distinct mechanical strains; for example, microridges are shorter and more branched in cells covering pectoral fins, and are reduced in periderm cells that stretch over bulges in the skin, such as those created by neuromast mechanosensory organs (not shown). This variation in microridge patterns provides an opportunity to further explore how actin regulators, contraction, and membrane biomechanics contribute to sculpting complex 3D membrane topographies.

## MATERIALS AND METHODS

### Zebrafish

Zebrafish (Danio rerio) were raised at 28.5C on a 14h/10h light/dark cycle. Embryos were raised at 28.5C in embryo water (0.3 g/L Instant Ocean Salt, 0.1% methylene blue). All experimental procedures were approved by the Chancellor’s Animal Research Care Committee at UCLA.

### Reagents

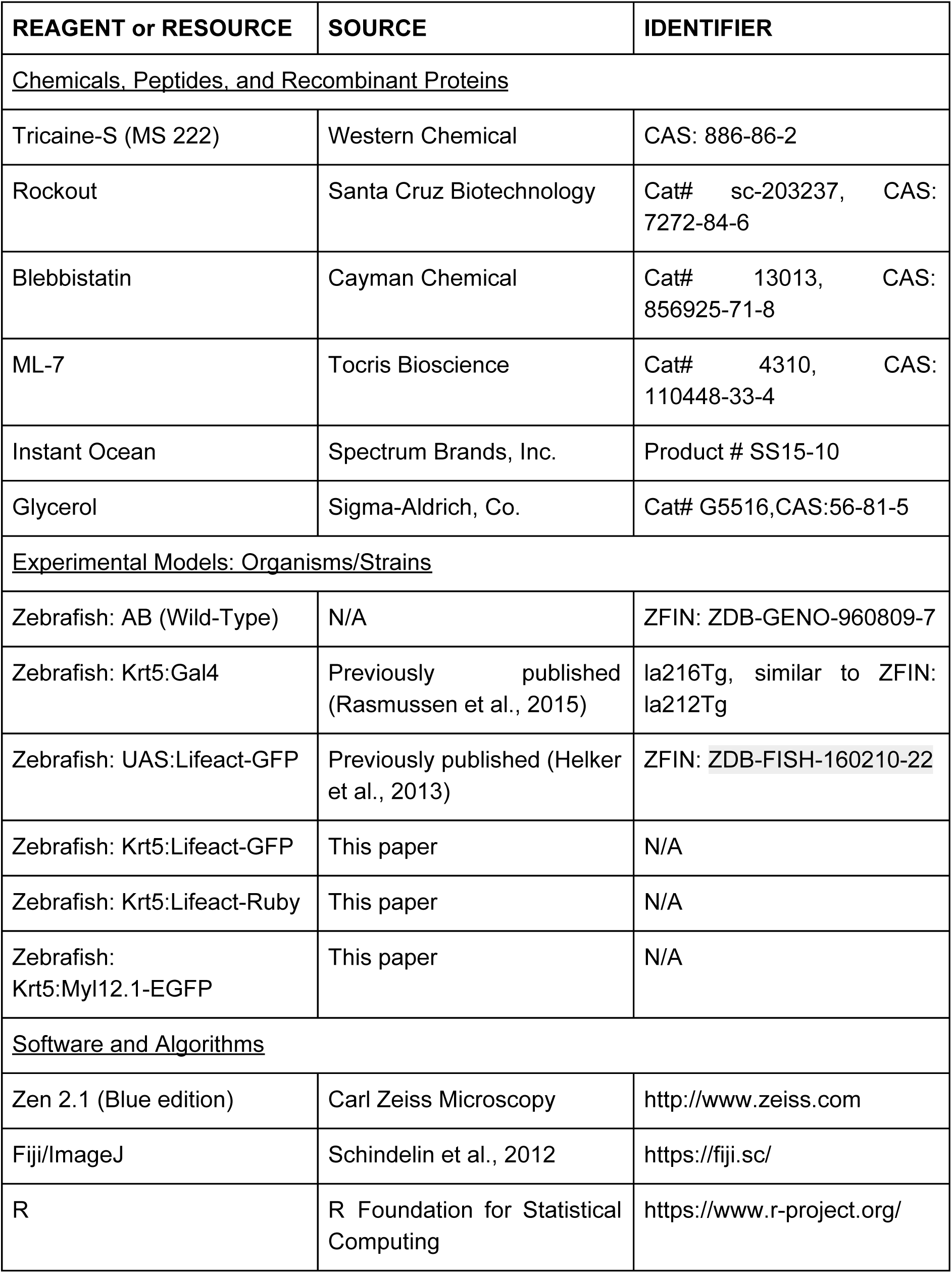
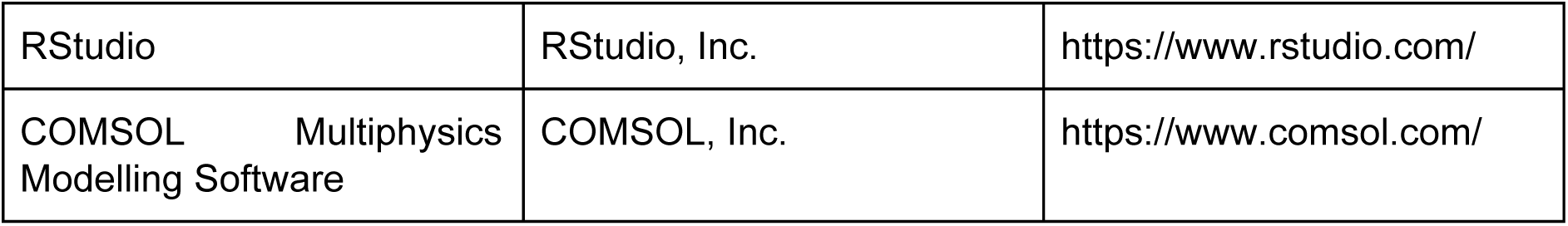

### Plasmids and Transgenes

Plasmids were constructed using the Gateway-based Tol2kit (Kwan et al., 2007). The following plasmids have been described previously: p5E-Krt5 (Rasmussen et al., 2015), pME-myl12.1 (Maître et al., 2012), p3E-polyA, p3E-EGFPpA, and pDestTol2pA2 (Kwan et al., 2007).

To construct pME-Lifeact-GFP, the following primers were used: (5’-GGGGACAAGTTTGTACAAAAAAGCAGGCTTAATGGGTGTCGCAGATTTG-3’, 5’-GGGGACCACTTTGTACAAGAAAGCTGGGTATTACTTGTACAGCTCGTC-3’; *actb1:lifeact-GFP*; (Behrndt et al., 2012));

To construct pME-Lifeact-Ruby, the following primers were used: (5’-GGGGACAAGTTTGTACAAAAAAGCAGGCTTAATGGGTGTCGCAGATTTG-3’, 5’-GGGGACCACTTTGTACAAGAAAGCTGGGTATTAAGCGCCTGTGCTATG-3’; *actb1:lifeact-RFP*; (Behrndt et al., 2012)).

### Live imaging of zebrafish embryos

Live zebrafish embryos were anaesthetized with 0.2mg/mL MS-222 in system water prior to mounting. Embryos were embedded in 1.2% agarose on a cover slip and sealed within a microscope chamber, as previously described (O’Brien et al., 2009). Chambers were filled with 0.2mg/mL MS-222 solution and sealed with vacuum grease.

### Periderm Cell Ablation

Periderm cells were ablated on a Zeiss LSM 880 microscope equipped with a 40x oil objective and a Coherent Chameleon Ultra II 2-photon laser set to 813nm, adapting of a previously described method (O’Brien et al., 2009). To ablate cells, the 488 laser was used to find and focus on the cell surface at 250x digital zoom. The cell was exposed to 2-photon laser illumination for 3-4 seconds at 5-6% laser power using “Live” scanning.

### Hyperosmolar Treatment

After mounting zebrafish embryos on coverslips in 1.2% agarose, slide chambers were filled with solutions of either 0.3g/mL Instant Ocean salt mix in DI water or 12.5% glycerol in Ringer’s Solution with 0.2 mg/mL MS-222. Time-lapse imaging was started immediately after filling the slide chamber with hyperosmolar media.

### Drug Treatments

All drugs were dissolved in DMSO. Treatment solutions were created by adding the appropriate volume of drug or an equivalent volume of DMSO (≤1%) to Ringer’s Solution with 0.2 mg/mL MS-222. Zebrafish larvae were immersed in treatment solutions for the specified period of time, then mounted in agarose and imaged while bathed in the same solution. For treatment periods longer than two hours, larvae were initially exposed to treatment solutions prepared without MS-222, then transferred to a treatment solution containing MS-222 at least 30 minutes prior to mounting and imaging.

### Microscopy

Live fluorescent imaging was performed on a Zeiss LSM 510, 800, or 880 confocal microscope using a 40x oil objective (1.3 NA) with 2-3x digital zoom.

### Image Analysis and Statistics

Image analysis was performed with FIJI (Schindelin et al., 2012). For display purposes, confocal z-stack images were projected (maximum intensity projection) and brightness and contrast were enhanced.

To analyze microridge length and other cell parameters, we developed an ImageJ script to automatically process cells in each image (Fig. S1). Cell outlines were traced by hand with the polygon selection tool to measure apical cell area. Brightness and contrast were automatically enhanced and the area around the cell was cleared. Images were then blurred using the Smoothen function three times, and passed through a Laplacian morphological filter from the MorphoLibJ plugin (Legland et al., 2016), using the square element and a radius of 1. Filtered images were automatically thresholded using the Triangle method and skeletonized. The Analyze Skeleton (2D/3D) feature was then used to measure microridge length. Statistical analyses and data presentation were conducted in RStudio.

To calculate surface excess in the case of pegs and microridges (Fig. 1G), we cropped 10×10 µm samples from the regions occupied by pegs or ridges in several cells and estimated the surface area of those samples with the following algorithm. We assumed that the height of the surface is proportional to the signal and that both pegs and microridges have the same maximal height of *h* = 400 nm. We normalized the samples, so that the 10th percentile became 0 and the 90th percentile 1, then used the following formula:

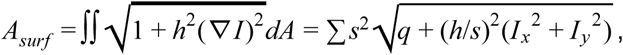

where ∇*I* is the gradient of the image (*I*_*x*_ and *I*_*y*_ are normalized sobel filters along x and y directions), *s* is the absolute pixel size and summation is taken over all pixels of the image. Surface excess of each sample was calculated as ε = *A*_*surf*_ /*A*_*proj*_ – 1, where *A*_*proj*_ is the areaof projected surface (width × length of the image).

To quantify NMII contraction, time-lapse z-stack images were projected and smoothened. Cell outlines were traced by hand and cells were cropped from each time-lapse frame. Brightness and contrast of each time-lapse frame was automatically adjusted, then images were automatically thresholded using the Triangle method. Thresholded pulses were measured using the ImageJ Analyze Particles function, excluding particles with < 4 pixels.

To construct the optic flow diagram (Fig. 4D’) we used FlowJ plugin in Fiji (Lucas & Kanade algorithm).

To study the angular distribution of surface structure coalescence (Fig. 7D), we analyzed videos from ablation experiments. We isolated distinct events of pegs coalescence and determined the direction (the line connecting pegs on a frame just before the fusion).

To calculate the direction and amount of elongation (Fig. 7F) we calculated the moment matrices M of cell shapes approximated with polygons (shifted to polygons’ centroids). Then we calculated non-rotational affine transformations that better transform moment matrix at frame n to moment matrix at frame n+1. The median direction of these transformations was chosen as the direction of elongation. The axes ratio was chosen as the elongation factor.

To calculate angular histograms for peg coalescence in the model (Fig. 8D,E), we created an automated version of the algorithm, which was used in the preparation of Figure 7D. This algorithm was applied to multiple (n = 100) simulations with random initial conditions and the averaged histograms plotted in Figures 8D,E.

To study how the angular distribution of peg coalescence in the model changes with the anisotropy parameter *ε* (Fig. 8F), we decreased the number of bins in the previous analysis to 2. All peg coalescence events with angles from −45° to 45° were considered as parallel to the axis of stretch, whereas those with angles from 45° to 135° as perpendicular to it. We then calculated the polarization parameter 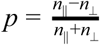, using the numbers of coalescence events in both bins. As defined, the value *p* =- 1 corresponds to all pegs merging perpendicular to the stretch axis, while *p* = 1 corresponds to all coalescence events being parallel to it.

### Modelling

We built a model as a system of partial differential equations and solved it with finite elements method. Calculations were performed with COMSOL Multiphysics (Burlington, MA, USA), postprocessing was done using custom python code.

#### Two layers

We subdivide the apical surface of the cell into two interacting subsystems. First layer is composite and represents the lipid membrane and the directly underlying it branched actin cytoskeleton, which fills out pegs and microridges. The second layer represents the actomyosin cortex. For simplicity, we assume that the interface between the two layers is flat and ignore its deformation (see below).

We assume that the two layers are mechanically coupled. Due to the presence of extensive cytoskeletal linkers connecting actomyosin to the transmembrane proteins embedded in the lipid bilayer, we introduce the no slip boundary condition. Thus, the two layers are coupled by the common strain field. This coupling has two important consequences. Firstly, velocity fields in the actomyosin layer that are induced by the forces directly translate onto the top layer and produce advection of the positioned there patterns, such as pegs or microridges. Secondly, we assume that contraction of actomyosin in the bottom layer reduces the tension in the top layer in a simple linear approximation.

The vertical component of interaction between the two layers results from the actin polymerization force in the top layer and is opposed by the oppositely directed force in the bottom layer (Gov, 2006). However, due to the differences in the layer thickness and, consequently, their mass, the effect of the actin polymerization force onto the actomyosin layer is assumed to be negligible (Mogilner, 2005). Thus, we ignore vertical forces and deformations of the bottom layer and assume that it remains flat.

#### Apical cortex layer

To describe actomyosin layer we used the active gel approach (Marchetti et al. 2013, Prost et al. 2015). Since we are interested in the behaviour on a very long timescale (minutes and hours), we can neglect inertia and shear elasticity and reduce the mechanics of the layer to a 2-dimensional compressible isotropic fluid with high viscosity and active stress. Although, in vivo cortical actomyosin is constantly assembled and disassembled, for the sake of simplicity, we assume the amount of actomyosin to be constant. Thus, we complement standard force balance equation (Prost et al, 2015):

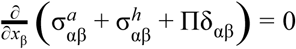

with the mass conservation law for the relative actomyosin density ρ:

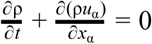

In the force balance equation, the indices α, β, γ∈{1, 2} refer to spatial coordinates *x, y* (*x*_1_, *x*_2_) in the plane of the layer, 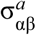 is the active stress, 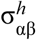 is the hydrodynamic (viscous) stress, and, finally, ∏ is the pressure. In all formulas repeating indices assume the Einstein summation convention and δ_αβ_ is the Kronecker delta. Hydrodynamic stress for a 2-dimensional fluid is defined as:

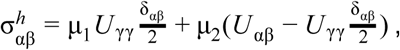

where *U* _αβ_ = ∂*u*_α_/∂*x*_β_ + ∂*u*_β_/∂*x*_α_ is a symmetric strain rate tensor, *u*_α_ is the velocity of the layer; μ_1_ and μ_2_ are the bulk and shear viscosities.

For constitutive relation we have used is a logarithmic continuation of the Hooke’s law: ∏ = ∏_0_ log ρ, where ρ is a dimensionless relative density of the layer and ∏_0_ is an effective stiffness of actomyosin.

#### Top composite layer

We describe the state of the top composite layer by a heuristic activator-inhibitor model of the Gierer-Meinhardt type (Gierer and Meinhardt, 1972) with two spatially distributed variables: the autocatalytic activator of actin polymerization *c*(*x, y*), and the height of the membrane measured relative to some initial plane *h*(*x, y*). This variable that plays the role of the inhibitor is presented in the Monge parametrization that is commonly used to describe the geometry of the membrane under the assumption | ∇*h*| ≪ 1 (Kabaso et al, 2011, Gov 2006).

Dynamics of activator *c* is represented by the following equation that directly follows the Gierer-Meinhardt form:

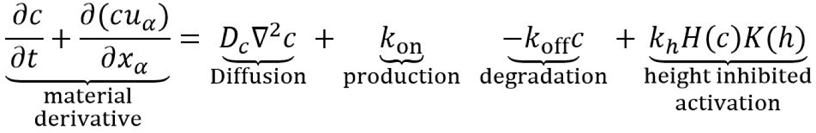

Parameters *D*_*c*_, *k*_*on*_, *k*_*off*_, *k*_*h*_, *c*_0_ and *h*_0_ are constants whose values are given in Table 1. The last term corresponds to the process of positive feedback with the rate that saturates when the concentration increases (Hill-like term) and diminishes when *h* increases:

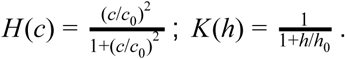

**Table 1.**
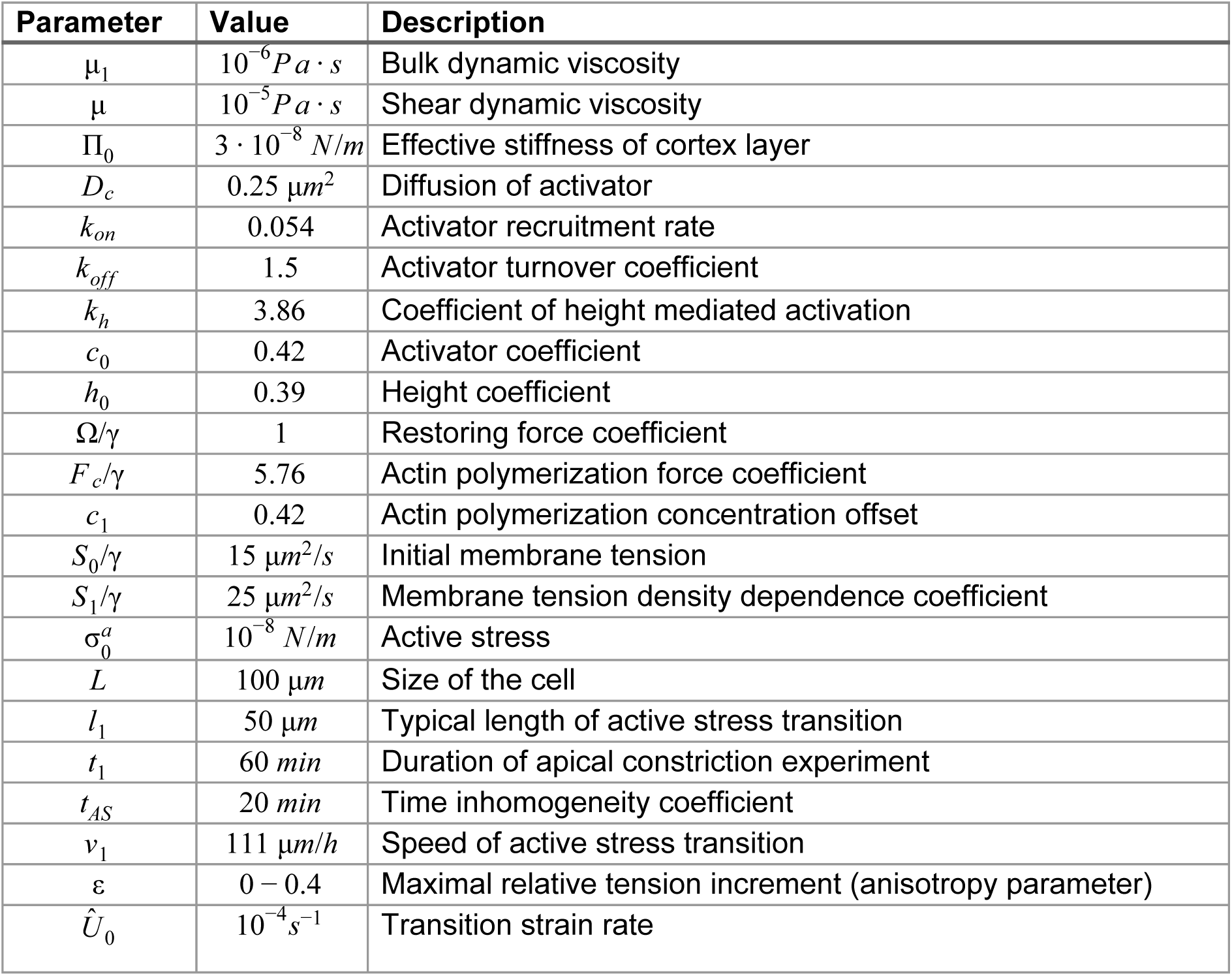
*of parameters*

Following (Gierer and Meinhardt, 1972), we chose the quadratic order in the Hill-like term as the smallest integer order which allows the system to be unstable and form spatial patterns.

We describe mechanical properties of the layer with the Helfrich Hamiltonian (Helfrich, 1973):

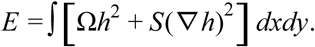

where *S* is the effective membrane tension coefficient. We also added a linear restoring force with coefficient Ω and neglected curvature-dependent energy terms. Minimizing elastic energy results in

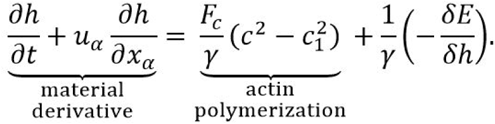

The actin polymerization force is postulated to depend quadratically on the concentration of activator *c* with the preferred concentration *c*_1_ and γ is the local Oseen parameter. The functional derivative has the form:

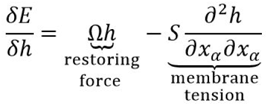

In the force equilibrium, the actin polymerization force is balanced by the restoring force and membrane tension.

Following from the earlier introduced mechanical coupling between the layers by strain, we posit that tension coefficient in the top composite layer depends on the relative density of the underlying actomyosin layer: when actomyosin contracts, the produced negative strain reduces the membrane layer tension. We chose a simple linear relation to reflect this:

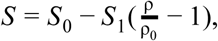

#### Apical constriction

To model pattern formation during the apical surface constriction, we applied time and space dependent active stress to a hexagonal cell with the side length *L*. We generated the initial pattern by simulating the equations for the lipid membrane layer with zero velocity field and pressure in cortex layer.

We first used spatially uniform active stress function:

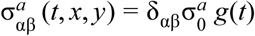

where 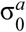 is the magnitude of active stress, and g(t) is a function representing temporal inhomogeneity: 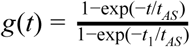, where *t*_*AS*_ is the time parameter, *t*_1_ is the time of simulation. To model the hypothesis that apical constriction initiates at the cell periphery, we introduced radially symmetric wave-like active stress function in the form:

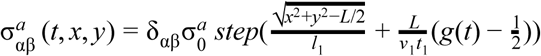

where 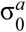 is the magnitude of active stress, *l*_1_ is the transition half-length, *v*_1_ is a typical wave propagation speed and *step*(*z*) is a continuous step function: 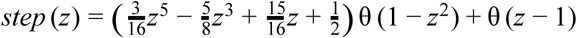, where θ (*z*) is Heaviside step function.

#### Anisotropic elongation

We speculate that during rapid uniaxial elongation of the cell, actomyosin filaments reorient in the direction of elongation (Fig. 8B) and thus introduce tensile anisotropy that propagates to the top layer. The membrane tension term becomes:

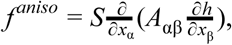

where *A*_αβ_ represents tensile anisotropic tensor which we chose as:

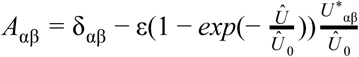

Here 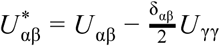 is the traceless component of the strain rate 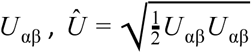 the positive eigenvalue of traceless matrix *U* _αβ_; ε is the maximal relative increment and is a typical strain-rate stress at which actomyosin filaments become partially ordered and _*Û*_ 0 membrane tension becomes sensitive to anisotropic flow. The idea behind this dependency is as follows. Without shear flow (*Û* _0_ = 0), the tension tensor is isotropic. With very large shear flow (*Û* _0_→∞), tension in the direction of elongation becomes smaller by factor 1 - ε and tension in the orthogonal direction becomes greater by factor 1 + ε. The transition occurs at a typical strain rate of *Û* _0_.

To simulate the elongation itself, we used a time-dependent affine transformation applied to the square border of the cell in order to simulate the experiment data (Fig. 7E,F). The duration of simulation was *t*_2_ = 15 *min*. We varied parameter ε from 0 (isotropic case, Video 6, left panel) to 0.4. Anisotropic case with ε = 0.25 is shown in Video 6 on the right panel.

### Online supplemental material

Fig. S1 shows the automatic image processing pipeline used to measure microridge length. Fig. S2 shows additional quantification methods and demonstrates that microridges lengthen as apical surfaces constrict during development. Fig. S3 shows time-lapse sequences illustrating three mechanisms of microridge growth. Fig. S4 illustrates the consequences of Arp2/3 inhibition on microridge length and apical constriction. Video 1 shows live imaging of microridge morphogenesis during development, demonstrating that microridges form from the accretion of pegs. Video 2 shows simulations of a biomechanical model demonstrating that reduction of membrane tension can promote centripetal micoridge development. Video 3 shows live imaging of an NMII reporter at 3 different developmental stages, revealing pulsatile apical contractions that pinch the membrane. Video 4 shows an *in vivo* live imaging experiment, demonstrating that reducing membrane tension with hyperosmolar media promotes microridge development. Video 5 shows a time-lapse sequence of a cell stretching after ablating neighboring cells. Video 6 shows simulations of a computational model demonstrating the effect of anisotropic cortical flow on the orientation of microridges during cell stretching.

## AUTHOR CONTRIBUTIONS

APvL, ISE, ABG, and AS conceived of and designed the study; APvL performed all zebrafish experiments; ISE, IVM, and ABG performed all theoretical experiments; APvL, ISE, ABG, and AS wrote the paper.

## ACKNOWLEDGMENTS

We thank members of the Sagasti lab and Margot Quinlan for comments on the work, and Son Giang for excellent fish care. We also appreciate image analysis advice from Dr. Roy Wollman and Dr. Alon Oyler-Yaniv. This work was funded by NIH grants R21EY024400 and R01GM122901 to AS and BBSRC grants BB/P01190X and BB/P006507 to ABG. APvL was supported by the Ruth L. Kirschstein National Research Service Award GM007185.

## VIDEO LEGENDS

**Video 1**

Live imaging of Lifeact-GFP in periderm cells during microridge development beginning at 18 hpf. All images are maximum intensity projections; time stamp represents hours:minutes:seconds.

**Video 2**

*In silico* simulation of apical constriction in our biomechanical model recapitulates the centripetal progression of microridge development observed *in vivo* (left panel). As the cell constricts its surface, membrane tension is relieved in a centripetally moving wave, promoting peg coalescence in a similar pattern. For comparison, right panel shows constriction in response to the spatially homogeneous increase in active stress.

**Video 3**

NMII (Myl12.1-EGFP) contractions in the apical cortex pull on actin microridges (Lifeact-Ruby) of periderm cells at various stages in microridge development, indicated by title cards. All images are maximum intensity projections; time stamp represents minutes:seconds.

**Video 4**

Live imaging of Lifeact-GFP in 16 hpf periderm cells, beginning 3 minutes after exposure to high salt media. All images are maximum intensity projections; time stamp represents minutes:seconds.

**Video 5**

Live imaging of Lifeact-GFP in 16 hpf periderm cells beginning immediately after ablation of periderm cells on opposite sides of the central cell. All images are maximum intensity projections; time stamp represents minutes:seconds.

**Video 6**

*In silico* simulation of uniaxial elongation of a model rectangular cell. Left panel shows pattern formation in the absence of sensitivity to anisotropic flow (ε = 0). Right panel shows pattern formation with membrane tension depending on underlying anisotropic actomyosin flow.

## REFERENCES

Atilgan, E., D. Wirtz, and S.X. Sun. 2006. Mechanics and dynamics of actin-driven thin membrane protrusions. Biophys. J. 90:65–76.

Behrndt, M., G. Salbreux, P. Campinho, R. Hauschild, F. Oswald, J. Roensch, S.W. Grill, and C.-P. Heisenberg. 2012. Forces driving epithelial spreading in zebrafish gastrulation. Science. 338:257–260.

Bereiter-Hahn, J., M. Osborn, K. Weber, and M. Vöth. 1979. Filament organization and formation of microridges at the surface of fish epidermis. J. Ultrastruct. Res. 69:316–330.

Bertet, C., L. Sulak, and T. Lecuit. 2004. Myosin-dependent junction remodelling controls planar cell intercalation and axis elongation. Nature. 429:667–671.

Blanchard, G.B., S. Murugesu, R.J. Adams, A. Martinez-Arias, and N. Gorfinkiel. 2010. Cytoskeletal dynamics and supracellular organisation of cell shape fluctuations during dorsal closure. Development. 137:2743–2752.

Blanchoin, L., R. Boujemaa-Paterski, C. Sykes, and J. Plastino. 2014. Actin dynamics, architecture, and mechanics in cell motility. Physiol. Rev. 94:235–263.

Blankenship, J.T., S.T. Backovic, J.S.P. Sanny, O. Weitz, and J.A. Zallen. 2006. Multicellular rosette formation links planar cell polarity to tissue morphogenesis. Dev. Cell. 11:459–470.

Bray, D., and J.G. White. 1988. Cortical flow in animal cells. Science. 239:883–888.

Buccione, R., J.D. Orth, and M.A. McNiven. 2004. Foot and mouth: podosomes, invadopodia and circular dorsal ruffles. Nat. Rev. Mol. Cell Biol. 5:647–657.

Cao, L.G., and Y.L. Wang. 1990. Mechanism of the formation of contractile ring in dividing cultured animal cells. II. Cortical movement of microinjected actin filaments. J. Cell Biol. 111:1905–1911.

David, D.J.V., A. Tishkina, and T.J.C. Harris. 2010. The PAR complex regulates pulsed actomyosin contractions during amnioserosa apical constriction in Drosophila. Development. 137:1645–1655.

DeBiasio, R.L., G.M. LaRocca, P.L. Post, and D.L. Taylor. 1996. Myosin II transport, organization, and phosphorylation: evidence for cortical flow/solation-contraction coupling during cytokinesis and cell locomotion. Mol. Biol. Cell. 7:1259–1282.

Depasquale, J.A. 2018. Actin Microridges: ACTIN MICRORIDGES IN EPITHELIUM. Anat. Rec. 31:81.

Derényi, I., F. Jülicher, and J. Prost. 2002. Formation and interaction of membrane tubes. Phys. Rev. Lett. 88:238101.

Fernández, B.G., A.M. Arias, and A. Jacinto. 2007. Dpp signalling orchestrates dorsal closure by regulating cell shape changes both in the amnioserosa and in the epidermis. Mech. Dev. 124:884–897.

Gierer, A., and H. Meinhardt. 1972. A theory of biological pattern formation. Kybernetik. 12:30–39.

Gorelik, J., A.I. Shevchuk, G.I. Frolenkov, I.A. Diakonov, M.J. Lab, C.J. Kros, G.P. Richardson, I. Vodyanoy, C.R.W. Edwards, D. Klenerman, and Y.E. Korchev. 2003. Dynamic assembly of surface structures in living cells. Proc. Natl. Acad. Sci. U. S. A. 100:5819–5822.

Gov, N.S. 2006. Dynamics and morphology of microvilli driven by actin polymerization. Phys. Rev. Lett. 97:018101.

Gov, N.S., and A. Gopinathan. 2006. Dynamics of membranes driven by actin polymerization. Biophys. J. 90:454–469.

Helfrich, W. 1973. Elastic properties of lipid bilayers: theory and possible experiments. Z. Naturforsch. C. 28:693–703.

Helker, C.S.M., A. Schuermann, T. Karpanen, D. Zeuschner, H.-G. Belting, M. Affolter, S. Schulte-Merker, and W. Herzog. 2013. The zebrafish common cardinal veins develop by a novel mechanism: lumen ensheathment. Development. 140:2776–2786.

Khaliullin, R.N., R.A. Green, L.Z. Shi, J.S. Gomez-Cavazos, M.W. Berns, A. Desai, and K. Oegema. 2018. A positive-feedback-based mechanism for constriction rate acceleration during cytokinesis in Caenorhabditis elegans. Elife. 7. doi: 10.7554/eLife.36073.

Kwan, K.M., E. Fujimoto, C. Grabher, B.D. Mangum, M.E. Hardy, D.S. Campbell, J.M. Parant, H.J. Yost, J.P. Kanki, and C.-B. Chien. 2007. The Tol2kit: a multisite gateway-based construction kit for Tol2 transposon transgenesis constructs. Dev. Dyn. 236:3088–3099.

Lam, P.-Y., S. Mangos, J.M. Green, J. Reiser, and A. Huttenlocher. 2015. In vivo imaging and characterization of actin microridges. PLoS One. 10:e0115639.

Legland, D., I. Arganda-Carreras, and P. Andrey. 2016. MorphoLibJ: integrated library and plugins for mathematical morphology with ImageJ. Bioinformatics. 32:3532–3534.

Lomakin, A.J., K.-C. Lee, S.J. Han, D.A. Bui, M. Davidson, A. Mogilner, and G. Danuser. 2015. Competition for actin between two distinct F-actin networks defines a bistable switch for cell polarization. Nat. Cell Biol. 17:1435–1445.

Maître, J.-L., H. Berthoumieux, S.F.G. Krens, G. Salbreux, F. Jülicher, E. Paluch, and C.-P. Heisenberg. 2012. Adhesion functions in cell sorting by mechanically coupling the cortices of adhering cells. Science. 338:253–256.

Marston, D.J., C.D. Higgins, K.A. Peters, T.D. Cupp, D.J. Dickinson, A.M. Pani, R.P. Moore, A.H. Cox, D.P. Kiehart, and B. Goldstein. 2016. MRCK-1 Drives Apical Constriction in C. elegans by Linking Developmental Patterning to Force Generation. Curr. Biol. 26:2079–2089.

Martin, A.C., and B. Goldstein. 2014. Apical constriction: themes and variations on a cellular mechanism driving morphogenesis. Development. 141:1987–1998.

Martin, A.C., M. Kaschube, and E.F. Wieschaus. 2009. Pulsed contractions of an actin-myosin network drive apical constriction. Nature. 457:495–499.

Mason, F.M., M. Tworoger, and A.C. Martin. 2013. Apical domain polarization localizes actin-myosin activity to drive ratchet-like apical constriction. Nat. Cell Biol. 15:926–936.

Matsumura, F. 2005. Regulation of myosin II during cytokinesis in higher eukaryotes. Trends Cell Biol. 15:371–377.

Mogilner, A., and B. Rubinstein. 2005. The physics of filopodial protrusion. Biophys. J. 89:782–795.

Munjal, A., J.-M. Philippe, E. Munro, and T. Lecuit. 2015. A self-organized biomechanical network drives shape changes during tissue morphogenesis. Nature. 524:351–355.

Nolen, B.J., N. Tomasevic, A. Russell, D.W. Pierce, Z. Jia, C.D. McCormick, J. Hartman, R. Sakowicz, and T.D. Pollard. 2009. Characterization of two classes of small molecule inhibitors of Arp2/3 complex. Nature. 460:1031–1034.

O’Brien, G.S., S. Rieger, S.M. Martin, A.M. Cavanaugh, C. Portera-Cailliau, and A. Sagasti. 2009. Two-photon axotomy and time-lapse confocal imaging in live zebrafish embryos. J. Vis. Exp. doi: 10.3791/1129.

Olson, K.R., and P.O. Fromm. 1973. A scanning electron microscopic study of secondary lamellae and chloride cells of rainbow trout (Salmo gairdneri). Z. Zellforsch. Mikrosk. Anat. 143:439–449.

Pinto, C.S., A. Khandekar, R. Bhavna, P. Kiesel, G. Pigino, and M. Sonawane. 2019. Microridges are apical epithelial projections formed of F-actin networks that organize the glycan layer. Sci. Rep. 9:12191.

Pollard, T.D. 2016. Actin and Actin-Binding Proteins. Cold Spring Harb. Perspect. Biol. 8. doi: 10.1101/cshperspect.a018226.

Pollard, T.D., and J.A. Cooper. 2009. Actin, a central player in cell shape and movement. Science. 326:1208–1212.

Prost, J., F. Jülicher, and J.-F. Joanny. 2015. Active gel physics. Nat. Phys. 11:111.

Raman, R., I. Damle, R. Rote, S. Banerjee, C. Dingare, and M. Sonawane. 2016. aPKC regulates apical localization of Lgl to restrict elongation of microridges in developing zebrafish epidermis. Nat. Commun. 7:11643.

Rasmussen, J.P., G.S. Sack, S.M. Martin, and A. Sagasti. 2015. Vertebrate epidermal cells are broad-specificity phagocytes that clear sensory axon debris. J. Neurosci. 35:559–570.

Ratheesh, A., G.A. Gomez, R. Priya, S. Verma, E.M. Kovacs, K. Jiang, N.H. Brown, A. Akhmanova, S.J. Stehbens, and A.S. Yap. 2012. Centralspindlin and α-catenin regulate Rho signalling at the epithelial zonula adherens. Nat. Cell Biol. 14:818–828.

Rauzi, M., P.-F. Lenne, and T. Lecuit. 2010. Planar polarized actomyosin contractile flows control epithelial junction remodelling. Nature. 468:1110–1114.

Roh-Johnson, M., G. Shemer, C.D. Higgins, J.H. McClellan, A.D. Werts, U.S. Tulu, L. Gao, E. Betzig, D.P. Kiehart, and B. Goldstein. 2012. Triggering a cell shape change by exploiting preexisting actomyosin contractions. Science. 335:1232–1235.

Rosenblatt, J., M.C. Raff, and L.P. Cramer. 2001. An epithelial cell destined for apoptosis signals its neighbors to extrude it by an actin- and myosin-dependent mechanism. Curr. Biol. 11:1847–1857.

Rotty, J.D., and J.E. Bear. 2014. Competition and collaboration between different actin assembly pathways allows for homeostatic control of the actin cytoskeleton. Bioarchitecture. 5:27–34.

Schindelin, J., I. Arganda-Carreras, E. Frise, V. Kaynig, M. Longair, T. Pietzsch, S. Preibisch, C. Rueden, S. Saalfeld, B. Schmid, J.-Y. Tinevez, D.J. White, V. Hartenstein, K. Eliceiri, P. Tomancak, and A. Cardona. 2012. Fiji: an open-source platform for biological-image analysis. Nat. Methods. 9:676–682.

Solon, J., A. Kaya-Copur, J. Colombelli, and D. Brunner. 2009. Pulsed forces timed by a ratchet-like mechanism drive directed tissue movement during dorsal closure. Cell. 137:1331–1342.

Sonal, J. Sidhaye, M. Phatak, S. Banerjee, A. Mulay, O. Deshpande, S. Bhide, T. Jacob, I. Gehring, C. Nuesslein-Volhard, and M. Sonawane. 2014. Myosin Vb mediated plasma membrane homeostasis regulates peridermal cell size and maintains tissue homeostasis in the zebrafish epidermis. PLoS Genet. 10:e1004614.

Sperry, D.G., and R.J. Wassersug. 1976. A proposed function for microridges on epithelial cells. Anat. Rec. 185:253–257.

Straight, A.F., A. Cheung, J. Limouze, I. Chen, N.J. Westwood, J.R. Sellers, and T.J. Mitchison. 2003. Dissecting temporal and spatial control of cytokinesis with a myosin II Inhibitor. Science. 299:1743–1747.

Straus, L.P. 1963. A Study of the Fine Structure of the So-Called Chloride Cell in the Gill of the Guppy Lebistes Reticulatus P. Physiol. Zool. 36:183–198.

Uehara, K., M. Miyoshi, and S. Miyoshi. 1988. Microridges of oral mucosal epithelium in carp, Cyprinus carpio. Cell Tissue Res. 251:547–553.

Uehara, K., M. Miyoshi, and S. Miyoshi. 1991. Cytoskeleton in microridges of the oral mucosal epithelium in the carp, Cyprinus carpio. Anat. Rec. 230:164–168.

Zihni, C., and S.J. Terry. 2015. RhoGTPase signalling at epithelial tight junctions: Bridging the GAP between polarity and cancer. Int. J. Biochem. Cell Biol. 64:120–125.

