## Supplemental Figures and Legends for "Cortical contraction drives the 3D patterning of epithelial cell surfaces"

SUPPLEMENTAL FIGURE LEGENDS

**Figure S1** Image analysis method for microridge detection
Stepwise illustration of image analysis pipeline, as described in Materials and Methods.

**Figure S2** Additional analyses of microridge development
(A) Box and violin plot of microridge length in periderm cells at the indicated stages of zebrafish development. For a weighted presentation of this data, see Fig. 1B. ***p < 0.0001; ANOVA followed by Tukey's HSD test (n=15582 structures in 23 cells from 10, 16 hpf fish; n=5096 structures in 40 cells from 9 fish at 24 hpf; n=4572 structures in 40 cells from 9 fish at 32 hpf; n=1309 structures in 19 cells from 6 fish at 48 hpf).
(B) Dot and box plot of average microridge length per periderm cell at the indicated stages of zebrafish development. ***p < 0.0001; ANOVA followed by Tukey's HSD test (n=15582 structures in 23 cells from 10 fish at 16 hpf; n=5096 structures in 40 cells from 9 fish at 24 hpf; n=4572 structures in 40 cells from 9 fish at 32 hpf; n=1309 structures in 19 cells from 6 fish at 48 hpf).
(C) Scatter plot of average microridge length per cell vs. apical cell area at the indicated stages of zebrafish development (n=15582 structures in 23 cells from 10 fish at 16 hpf; n=5096 structures in 40 cells from 9 fish at 24 hpf; n=4572 structures in 40 cells from 9 fish at 32 hpf; n=1309 structures in 19 cells from 6 fish at 48 hpf).

**Figure S3** Three modes of micoridge formation and growth
(A-C) Time-lapse sequences show examples of the coalescence of two pegs to form an incipient microridge (A), the addition of a peg to the end of a pre-existing microridge (B), and the joining of two pre-existing microridges (C).
Scale bars: 1μm (A)

**Figure S4** Arp2/3 activity is required for microridge development, but not apical constriction
(A) Representative projections of Lifeact-GFP in periderm cells on 24 hpf zebrafish larvae after 8 hr treatment with either 1% DMSO or 100μM CK666.
(B) Box and violin plot of microridge length in 24 hpf zebrafish larvae after 8 hr treatment with either 1% DMSO or 100μM CK666. Data displayed is a weighted distribution of microridge length where frequency is proportional to microridge length, approximating occupied area. ***p < 0.0001; student's t-test (n=5283 structures in 44 cells from 11 fish for 1% DMSO, n=6130 structures in 40 cells from 12 fish for 100μM CK666).
(C) Dot and box plot of periderm cell apical area in 24 hpf zebrafish embryos after 8 hr treatment with either 1% DMSO or 100μM CK666. ***p < 0.0001; student's t-test (n=44 cells from 11 fish for 1% DMSO, n=40 cells from 12 fish for 100μM CK666).
Scale bars: 10μm (A)

### Figure S1

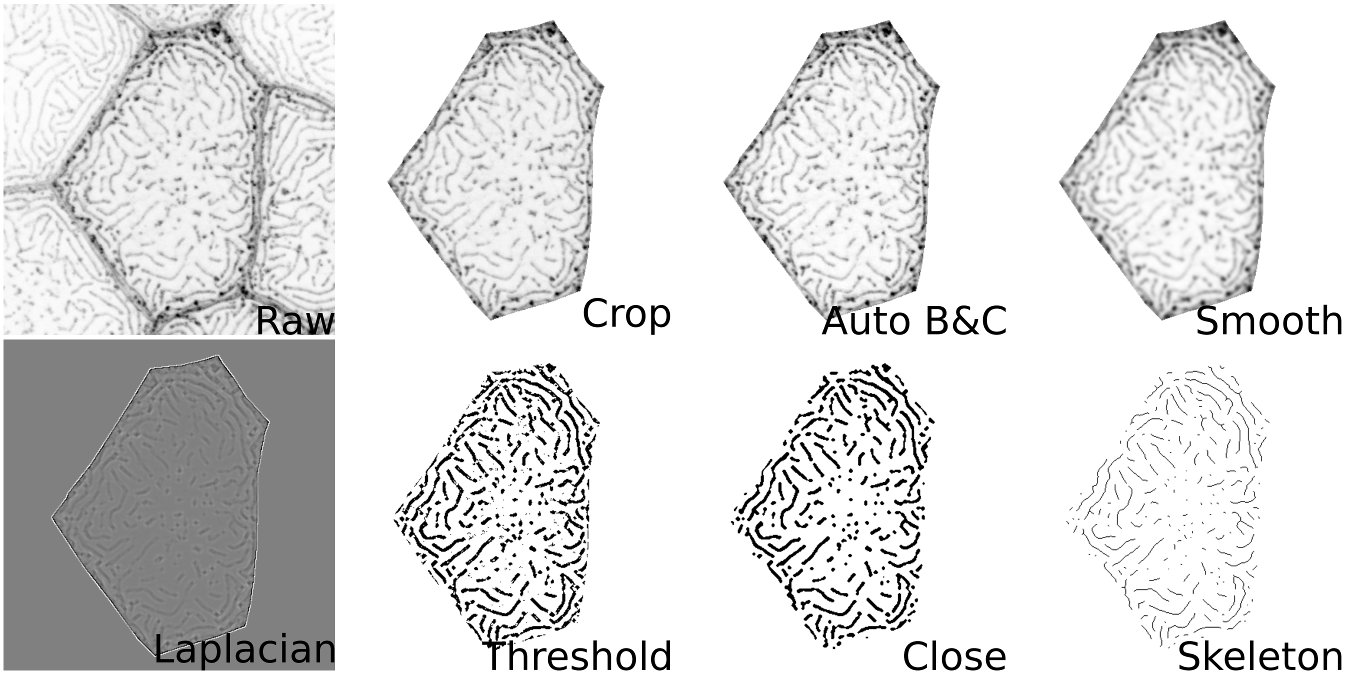
Raw
Crop
Auto B&C
Smooth
Laplacian
Threshold
Close
Skeleton

### Figure S2

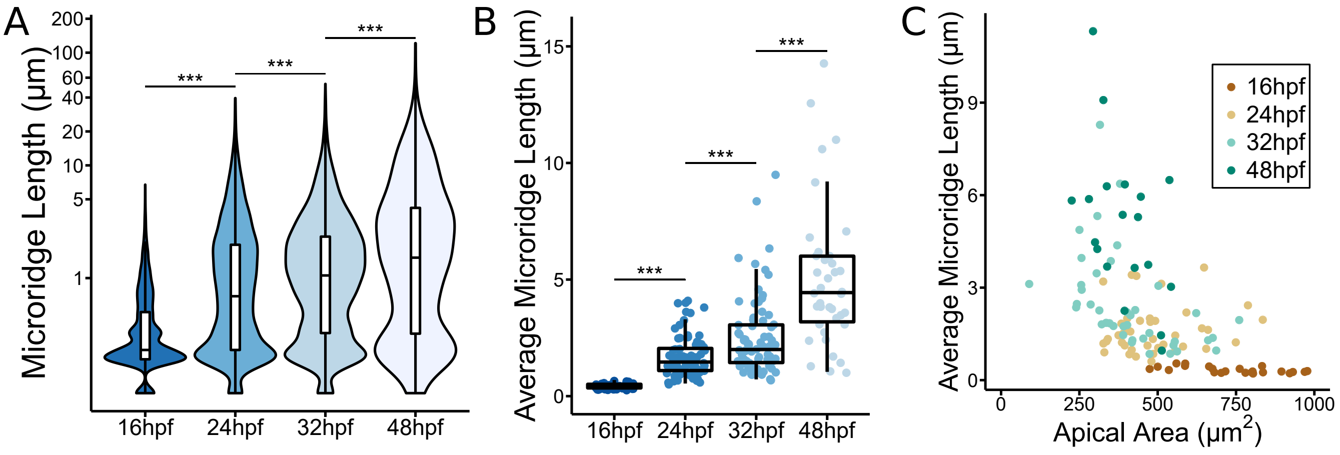
A
Microridge Length (µm)
200
100
60
40
20
10
5
1
***
***
***
16hpf
24hpf
32hpf
48hpf
B
Average Microridge Length (µm)
15
10
5
0
***
***
***
16hpf
24hpf
32hpf
48hpf
C
Average Microridge Length (µm)
9
6
3
0
0
250
500
750
1000
Apical Area (µm$^2$)
16hpf
24hpf
32hpf
48hpf

### Figure S3

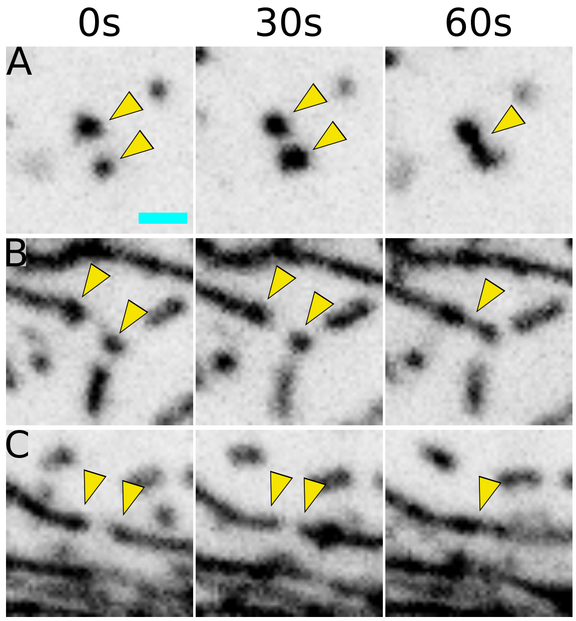

### Figure S4

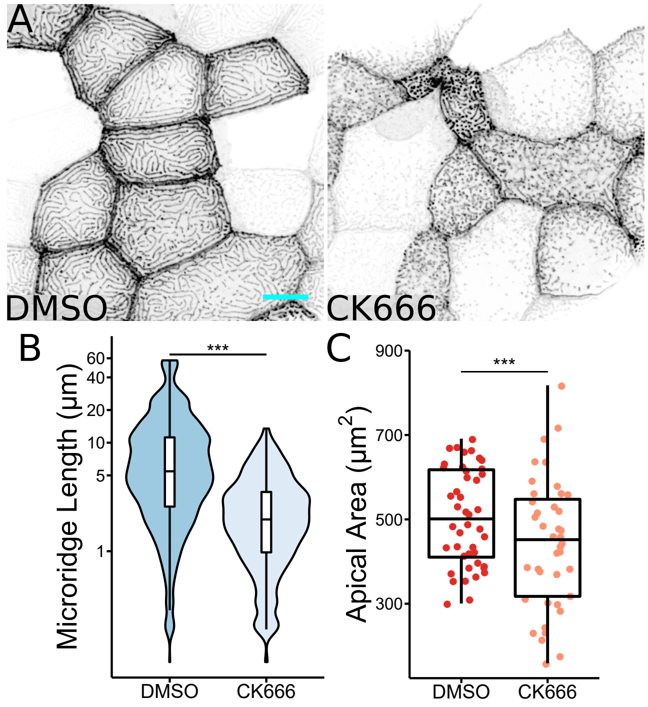

A
DMSO
CK666
B
***
Microridge Length (µm)
60
40
20
10
5
1
DMSO
CK666
C
***
Apical Area (µm2)
900
700
500
300
DMSO
CK666
